# Egr1 deficiency induces browning of inguinal subcutaneous white adipose tissue in mice

**DOI:** 10.1101/150003

**Authors:** Cécile Milet, Marianne Bléher, Kassandra Allbright, Mickael Orgeur, Fanny Coulpier, Delphine Duprez, Emmanuelle Havis

**Affiliations:** Sorbonne Universités, UPMC Univ Paris 06, CNRS UMR7622, Inserm U1156, IBPS-Developmental Biology Laboratory, F-75005 Paris, France.; University of Pittsburgh, Pittsburgh, Pennsylvania, United States.; École normale supérieure, PSL Research University,CNRS, Inserm,Institut de Biologie de l’École normale supérieure (IBENS), Plateforme Génomique, 75005 Paris, France.

**Keywords:** Egr1, Ucp1, transcription, matrix, browning, adipocyte, mouse

## Abstract

Beige adipocyte differentiation within white adipose tissue, referred to as browning, is seen as a possible mechanism for increasing energy expenditure. The molecular regulation underlying the thermogenic browning process has not been entirely elucidated. Here, we identify the zinc finger transcription factor EGR1 as a negative regulator of the beige fat program. Loss of *Egr1* in mice promotes browning in the absence of external stimulation and activates *Ucp1* that encodes the key thermogenic mitochondrial uncoupling protein-1. Moreover, EGR1 is recruited to the proximal region of the *Ucp1* promoter in subcutaneous inguinal white adipose tissue. Transcriptomic analysis of subcutaneous inguinal white adipose tissue in the absence of *Egr1* identifies the molecular signature of white adipocyte browning downstream of *Egr1* deletion and highlights a concomitant increase of beige differentiation marker and decrease in extracellular matrix gene expression. Conversely, *Egr1* overexpression in mesenchymal stem cells decreases beige adipocyte differentiation, while increasing extracellular matrix production. These results uncover the role of *Egr1* in blocking energy expenditure via direct *Ucp1* transcription regulation and highlight *Egr1* as a therapeutic target for counteracting obesity.

## Introduction

White fat browning is a mechanism that produces heat and limits weight gain. The understanding of the molecular regulation underlying white fat browning has sparked interest to counteract obesity.

The adipose tissue of humans and other mammals contains white adipose tissue (WAT) and brown adipose tissue (BAT). WAT and BAT are developmentally and functionally distinct and contain white and brown adipocytes, respectively (Berry et al., 2013; Bartelt and Heeren 2014; Harms and Seale, 2013). More recently, a third type of adipocytes has been described within WAT, beige adipocytes. Morphological and molecular analyses showed that brown and beige adipocytes are remarkably similar and express the same thermogenic markers (Kajimura et al., 2015). However beige adipocytes, in contrast to brown adipocytes, express thermogenic markers only after external stimulations, such as cold exposure, starvation, exercise or hormone treatment (Rosenwald et al., 2013). In the adult, beige adipocytes are produced by the trans-differentiation of mature white adipocytes (Kajimura et al., 2015) or by *de novo* differentiation of progenitors (Wang et al., 2013) in response to external stimulations. This process is referred to as “browning” or “beigeing” (Bartelt and Heeren, 2014, Garcia et al., 2016).

Because the increase of WAT is observed in many metabolic diseases, WAT browning represents a promising therapeutic approach. Consequently, it is crucial to decipher the molecular aspects underlying the beige differentiation program. Adipogenesis is triggered by a common adipogenic network, starting with the expression of *Cebpb* (CCAAT/enhancer binding protein ß), which activates the expression of *Pparg (*Peroxisome proliferator-activated receptor γ) and *Cebpa* (CCAAT/enhancer binding protein α), which in turn activates *Ppara* (Peroxisome proliferator-activated receptor α) expression (Peirce et al., 2014). Consistent with it is thermogenic function, brown/beige differentiated adipocytes express high levels of UCP1, a mitochondrial protein that uncouples oxidative phosphorylation from ATP synthesis (Klaus et al., 1991; Shabalina et al., 2013). The Krebs cycle enzymes, such as OGDH (oxoglutarate dehydrogenase), SUCLA2 (succinate-Coenzyme A ligase) and COX8B (Cytochrome C Oxidase Subunit VIIIb) (Forner et al., 2009; Wu et al., 2012) are also involved in heat production in beige/brown adipose tissue. Consistent with their anti-fat function, brown/beige differentiated adipocytes express factors involved in lipolysis such as PLIN5 (Perilipin 5; Gallardo-Montejano et al., 2016) and CIDEA (Cell Death-Inducing DFFA-Like Effector A; Wu et al., 2012). Beige adipocyte differentiation relies on the expression of a set of transcriptional activators (Bartelt and Heeren 2014; Harms and Seale 2013). PRDM16 (PR domain containing 16) is considered as a master regulator of the brown/beige program via direct interaction with transcription factors, such as C/EBPβ PPARα, PPARγ, and PGC-1α (Peroxisome proliferator-activated receptor Gamma Coactivator 1-alpha) (Rajakumari et al., 2013; Seale et al., 2011; Puigserver et al., 1998). Of note, beige and white differentiation programs share transcriptional regulators, such as C/EBPβ which has been shown to be sufficient for *Ucp1* transcription via direct binding to *Ucp1* proximal promoter in vitro (Yubero et al., 1994; reviewed in Villarroya et al., 2016). Moreover, *Cebpb* mutant mice display defective thermoregulation (Carmona et al., 2005). In addition to transcriptional regulators, growth factors such as FGF21 (Fibroblast Growth Factor-21) and BMP4 (Bone morphogenetic Protein-4), adipokines such as leptin and hormones such as T_3_ (Triiodothyronin 3) have been identified as being able to induce the brown/beige fat phenotype (Bartelt and Heeren, 2014; Kim and Plutzky, 2016; Forest et al., 2016). The T_4_ to T_3_ converting enzyme Desiodase 2 (DIO2) is also involved in the browning process (De Jesus et al., 2001).

The zinc finger transcription factor EGR1 (Early Growth Response-1) is involved in multiple processes including cell proliferation, differentiation, migration, apoptosis, and inflammation (Beckmann and Wilce, 1997; Cao et al., 1993; Pagel and Deindl, 2011; Sakamoto et al., 1994; Tsai-Morris et al., 1988) in many cell types. *Egr1* is expressed in adult adipose tissues (Yu et al., 2011; Zhang et al., 2013), where its overexpression has been linked to obesity in both humans and mouse models (Yu et al., 2011; Zhang et al., 2013). Consistently, EGR1 inhibits lipolysis and promotes fat accumulation in cultured adipocytes by directly repressing the transcription of the adipose triglyceride lipase (ATGL) gene (Chakrabarti et al., 2013).

In this study, we analysed the consequences of *Egr1* inhibition for subcutaneous inguinal white adipose tissue (SC-WAT) formation during postnatal and adult periods, using a mouse model deficient for *Egr1,* with no external stimulation. We also assessed the consequences of *Egr1* overexpression for beige differentiation in mesenchymal stem cells.

## Results and Discussion

### *Egr1^-/-^* mice display inguinal subcutaneous white adipose tissue browning with no external stimulation

The subcutaneous inguinal white adipose tissue (SC-WAT) expands during the post-natal period (Cereijo et al., 2014) and is the largest white fat depot in mice (Shabalina et al., 2013; Waldén et al., 2012). *Egr1* expression in SC-WAT was detected in blood vessels (Figure 1D, arrow a) as previously described (Khachigian et al., 1996) and in white adipocytes (Figure 1D, arrows b,c). The weight of SC-WAT fat pads was similar in *Egr1*^*+/+*^ and *Egr1*^*-/-*^ 4-month-old mice, although the total body weight was slightly reduced in *Egr1*^*-/-*^mice compared to control mice (Figure 1A-C). SC-WAT from 1-month-old (post-natal) and 4-month-old (adult) *Egr1*^*-/-*^mice exhibited high number of beige adipocytes, labelled with UCP1, compared to *Egr1*^*+/+*^mice (Figure 1E,F). Consistently, the *Ucp1* mRNA expression levels were increased in *Egr1*-deficient SC-WAT compared to equivalent control SC-WAT (Figure 1H). The density of adipocytes increased in SC-WAT of *Egr1*^*-/-*^mice compared to *Egr1*^*+/+*^mice with a significant increase of the proportion of beige adipocytes and thus a reduction in the proportion of white adipocytes (Figure 1G). Cell counts are compatible with two mechanisms for WAT browning described in the literature (Kajimura et al., 2015, Wang et al., 2013), trans-differentiation of white adipocytes and proliferation of beige cells. The increase of *Ucp1* transcript levels, of UCP1 protein and in the density of UCP1 + cells in SC-WAT of *Egr1*^*-/-*^mice (Figure 1) is consistent with the UCP1 increase previously observed in *Egr1*^-/-^mice under high fat diet feeding (Zhang et al., 2013). It has to be noted that there was no need of any high fat diet to observe *Ucp1*/UCP1 increase in SC-WAT fat pads of our *Egr1*^*-/-*^mice. We conclude that *Egr1* deficiency promotes spontaneous WAT browning without external stimulation. These results indicate that the presence of *Egr1* in white adipocytes represses WAT browning.

**Figure 1.**
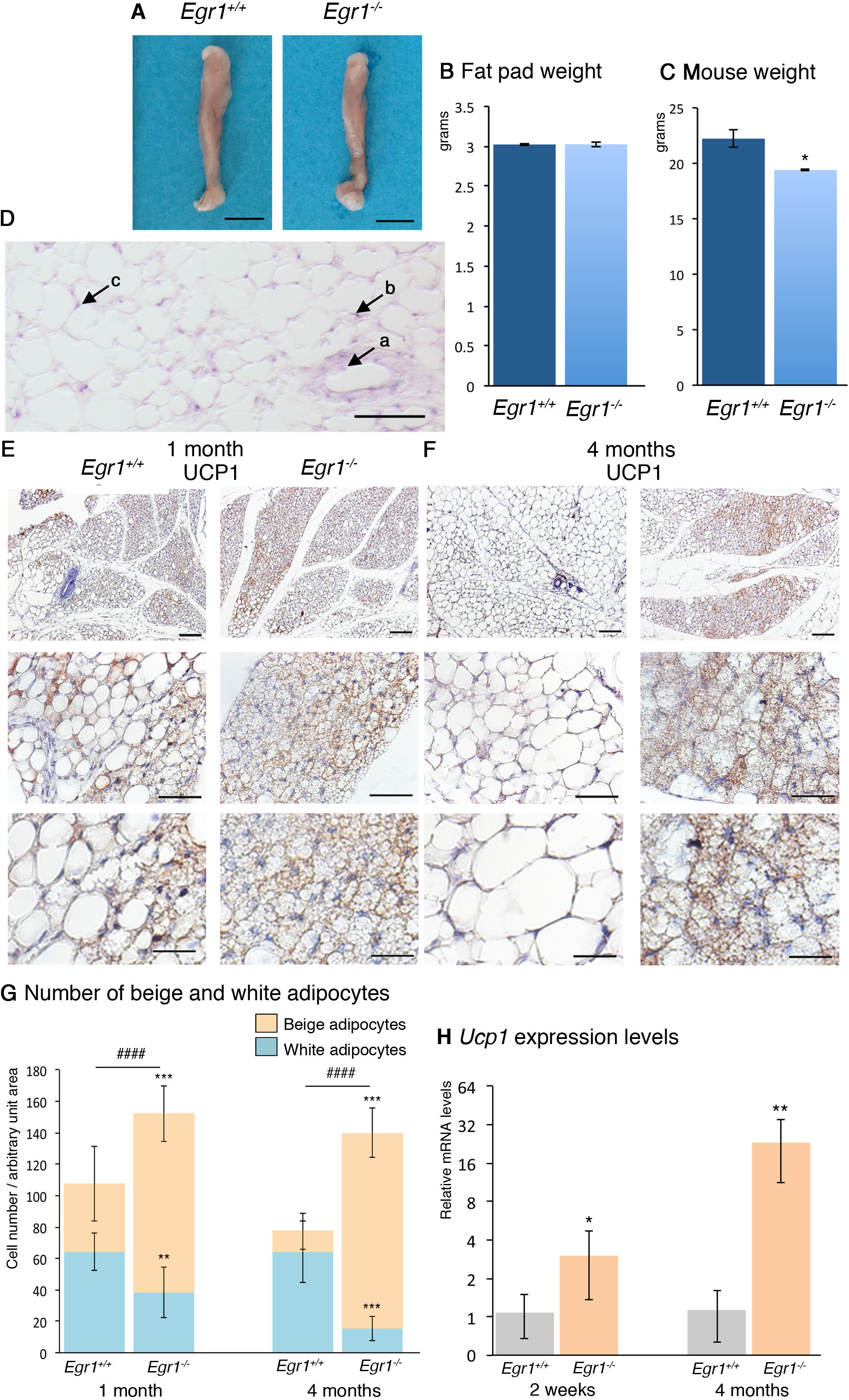
*Egr1* loss-of-function leads to inguinal subcutaneous white adipose tissue browning in postnatal and 4 month-old mice. (**A**) Pictures of fat pads (SC-WAT) from 4-month-old *Egr1*^*+/+*^and *Egr1*^*-/-*^mice. Scale bars: 5mm. (**B**) Weight in grams of SC-WAT of 4-month-old *Egr1*^*+/+*^and *Egr1*^*-/-*^mice. The graph shows mean ± standard deviations of 6 *Egr1*^*+/+*^fat pads and 8 *Egr1*^*-/-*^fat pads. (**C**) Weight in grams of 4-month-old wild-type and mutant mice. The graph shows means ± standard deviations of 4 *Egr1*^*+/+*^and 4 *Egr1*^*-/-*^mice. The p-value was obtained using the Mann-Whitney test. Asterisk indicates the p-value *P<0.05. (**D**) SC-WAT of 1-month-old mouse was longitudinally sectioned. 6μm sections were hybridized with the DIG-labeled antisense probe for *Egr1* (blue). Arrow “a” points *Egr1* expression in blood vessels. Arrows “b” and “c” indicate *Egr1* expression in white adipocytes. Scale bars: 50μm. (**E,F**) Sections of SC-WAT of 1-month-old (**E**) and 4-month-old (**F**) *Egr1*^*+/+*^and *Egr1*^*-/-*^mice were immuno-stained with UCP1 antibody and counterstained with hematoxilin. Scale bars: lower magnification 100μm; intermediate magnification 50μm, higher magnification 25μm. (**G**) White and beige adipocyte number was counted in arbitrary unit areas of transverse sections of the SC-WAT of 1 month-old (N = 7) and 4 month-old (N = 8) *Egr1*^*+/+*^and *Egr1*^*-/-*^mice. Graphs show means of 7 or 8 sections for each sample ± standard deviations. Asterisks indicate the p-values obtained using the Mann-Whitney test, comparing beige or white adipocyte number between mutant and control mice *** P<0.001, **P<0.01. Number signs indicate the p-values obtained using Anova test comparing cell number between mutant and control mice ### P<0.001. (**H**) RT-qPCR analysis of expression levels for beige adipocyte differentiation marker *Ucp1* in SC-WAT of 2-week-old and 4-month-old *Egr1*^*-/-*^mice compared to *Egr1*^*+/+*^mice. The mRNA levels of control (*Egr1*^*+/+*^) SC-WAT were normalized to 1. Graphs show means ± standard deviations of 7 samples from 2-week-old *Egr1*^*+/+*^mice, 5 samples from 2-week-old *Egr1*^*-/-*^mice and 5 samples from 4-month-old wild-type and mutant mice. The p-values were obtained using the Mann-Withney test. Asterisks indicate the p-values * p<0.05, ** p<0.01.

### Molecular signature of inguinal subcutaneous white adipose tissue browning downstream of *Egr1*

In order to define the molecular signature underlying WAT browning downstream of Egr1, we performed RNA-sequencing of SC-WAT of 2-week-old *Egr1*^*+/+*^and *Egr1*^*-/-*^mice. 336 differentially expressed genes were significantly detected in *Egr1*-deficient SC-WAT compared to control SC-WAT. The 132 upregulated differentially expressed genes (Figure 2A, Figure 2-figure supplement 1) were subjected to functional annotation clustering according to their Gene Ontology (GO) classification, in the “*Biological Process*” category (Figure 2-figure supplement 2). Among the 132 upregulated genes, the GO terms “*NADH metabolic process*”, “*Tricarboxylic acid cycle*”, “*Brown fat cell differentiation*” and “*Fatty acid metabolic process*” exhibited the highest enrichment scores (Figure 2-figure supplement 2). Consistent with the beige phenotype (Figure 1), the key beige adipocyte markers, *Ppargc1a, Ucp1*, *Cox8b*, *Cidea* (Garcia et al., 2016) and other genes known to be involved in the beige differentiation program, *Dio2*, *Pank1*, *Plin5*, *Ogdh* and *Sucla2* (De Jesus et al., 2001; Christian, 2014; Rosell et al., 2014; Forner et al., 2009) were identified as upregulated genes (Figure 2A). The increased expression of these beige genes was confirmed by RT-qPCR at 2 weeks and 4 months (Figure 2B,C). In addition, the generic adipogenesis regulators also known to be involved in beige differentiation, *Cepbb* (Kajimura et al., 2009) and *Ppara* (Barberá et al., 2001) displayed an increased expression in *Egr1*-deficient SC-WAT (Figure 2A-C). Interestingly, there was no modification of expression of signalling molecules controlling beige differentiation such as FGF21, BMP4 or Leptin. This indicates that EGR1 negatively regulates the transcription of beige differentiation markers. To test whether this regulation was direct, we performed Chromatin immunoprecipitation (ChIP) experiments from the SC-WAT of 2-week-old mice on key beige markers. EGR1 was recruited to the *Ucp1* proximal promoter in SC-WAT (Figure 2D), showing a direct transcriptional regulation by EGR1. EGR1 was also recruited to the *Cebpb* promoter but not to that of *Ppapgc1* gene (Figure 2D), highlighting a direct and an indirect transcriptional regulation of these two genes by EGR1. These results show that EGR1 exerts its transcriptional repression of the beige program at two levels at least, through the direct recruitment of the main beige differentiation marker *Ucp1* and also through the direct recruitment to the *Cebpb* gene, which is known to regulate *Ucp1* transcription (Yubero et al., 1994).

**Figure 2.**
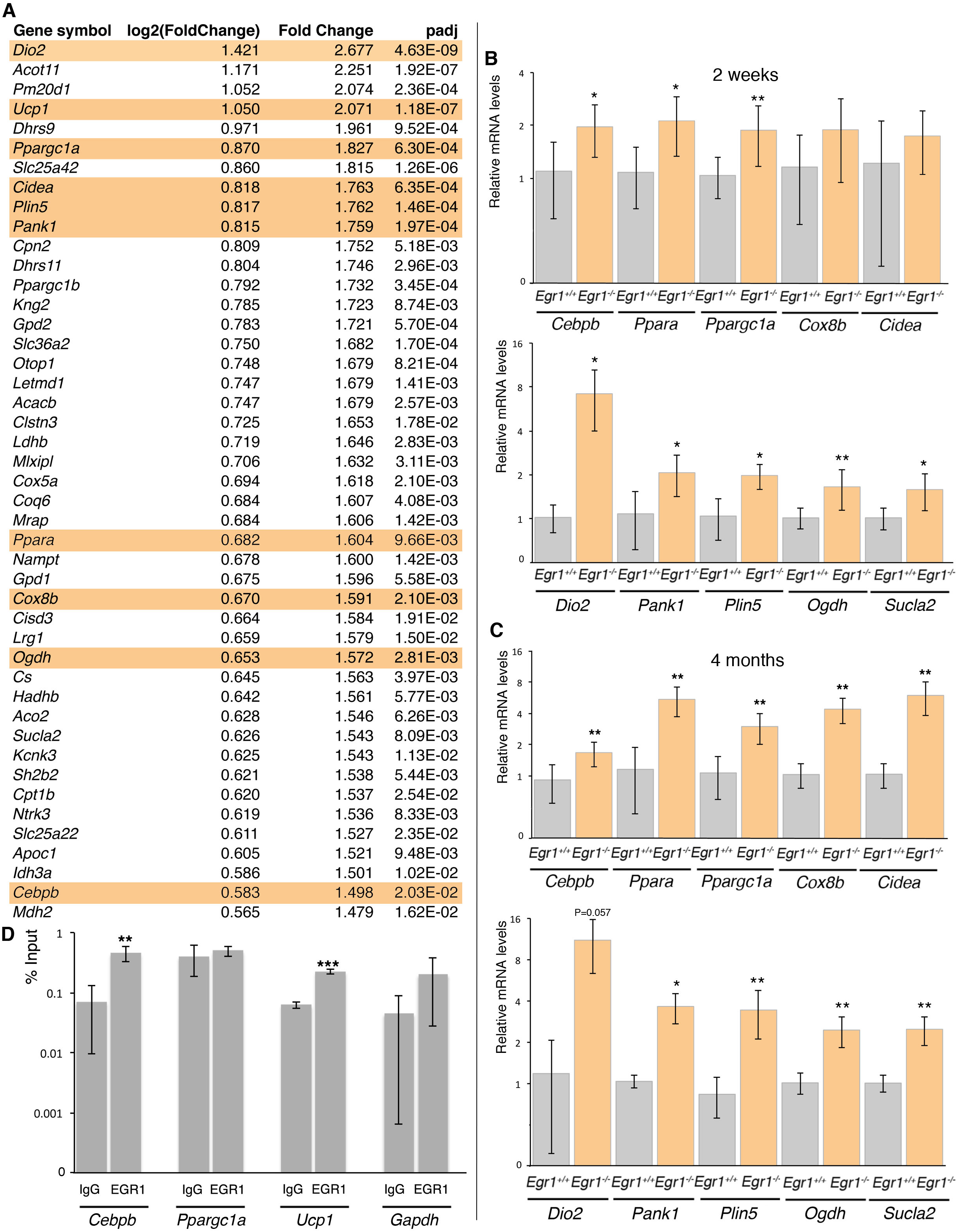
Transcriptomic analysis of subcutaneous inguinal adipose tissue of postnatal *Egr1*^*-/-*^versus *Egr1*^*+/+*^ mice shows upregulation of beige adipocyte markers. (**A**) List of the first 45 upregulated genes in 3 samples of SC-WAT of *Egr1*^*-/-*^versus *Egr1*^*+/+*^2-week-old mice. (**B,C**) RT-qPCR analysis of the expression levels for generic adipocyte differentiation markers *Cebpb*, *Ppara*, beige adipocyte differentiation marker, *Ppargc1a*, *Cox8b*, *Cidea, Dio2*, *Pank1, Plin5, Ogdh* and *Sucla2* in SC-WAT of 2-week-old (**B**) and 4-month-old (**C**) *Egr1*^*-/-*^mice compared to *Egr1*^*+/+*^mice. For each gene, the mRNA levels of control (*Egr1*^*+/+*^) SC-WAT were normalized to 1. Graphs show means ± standard deviations of 7 samples from 2-week-old *Egr1*^*+/+*^mice, 5 samples from 2-week-old *Egr1*^*-/-*^mice, and 5 samples from 4-month-old wild type and mutant mice. The p-values were obtained using the Mann-Withney test. Asterisks indicate the p-values * p<0.05, ** p<0.01. (**D**) ChIP assays were performed from 20 fat pads of 2-week-old mice with antibodies against EGR1 or IgG2 as irrelevant antibody in three independent biological experiments. ChIP products were analyzed by RT-q-PCR (N = 2). Primers targeting the proximal promoter regions of *Cebpb* and *Ucp1* revealed the recruitment of EGR1 in the vicinity of these sequences, while primers targeting the proximal promoter regions of *Ppargc1a* and *Gapdh* (negative controls) did not show any immunoprecipitation with EGR1 antibody compared to IgG2 antibody. Results were represented as percentage of the input. Error bars showed standard deviations. The p-values were obtained using the Mann-Withney test. Asterisks indicate the p-values, ** p<0.01, *** p<0.001.

The 204 downregulated differentially expressed genes (Figure 3A, Figure 3 – figure supplement 1) in SC-WAT of *Egr1*^*-/-*^mice were enriched for the GO terms *“Collagen fibril organization*”, “*Collagen catabolic process*” and “*Extracellular matrix organization*” (Figure 3-figure supplement 2). WAT produces extracellular matrix (ECM) whose composition and remodelling is crucial for adipocyte function (Mariman and Wang, 2010). Conversely, the expansion of adipose tissue during obesity leads to tissue remodelling and is associated with overexpression of *Col1a1*, *Col5a2*, *Fn1*, *Dcn* and the matrix metalloprotease *Mmp2* genes (Divoux and Clement, 2011; Berger et al., 2015; Bolton et al., 2008; Dubois et al., 2008; Henegar et al., 2008). In the transcriptome of *Egr1*-deficient SC-WAT, *Col1a1*, *Col1a2*, *Col5a2*, *Col14a1*, *Fn1*, *Post*, *Dcn* and *Mmp2* were downregulated (Figure 3A), which was confirmed by RT-qPCR in SC-WAT of 2 week-and 4 month-old mice (Figures 3B,C). We conclude that *Egr1*-deficiency represses ECM genes associated with obesity.

**Figure 3.**
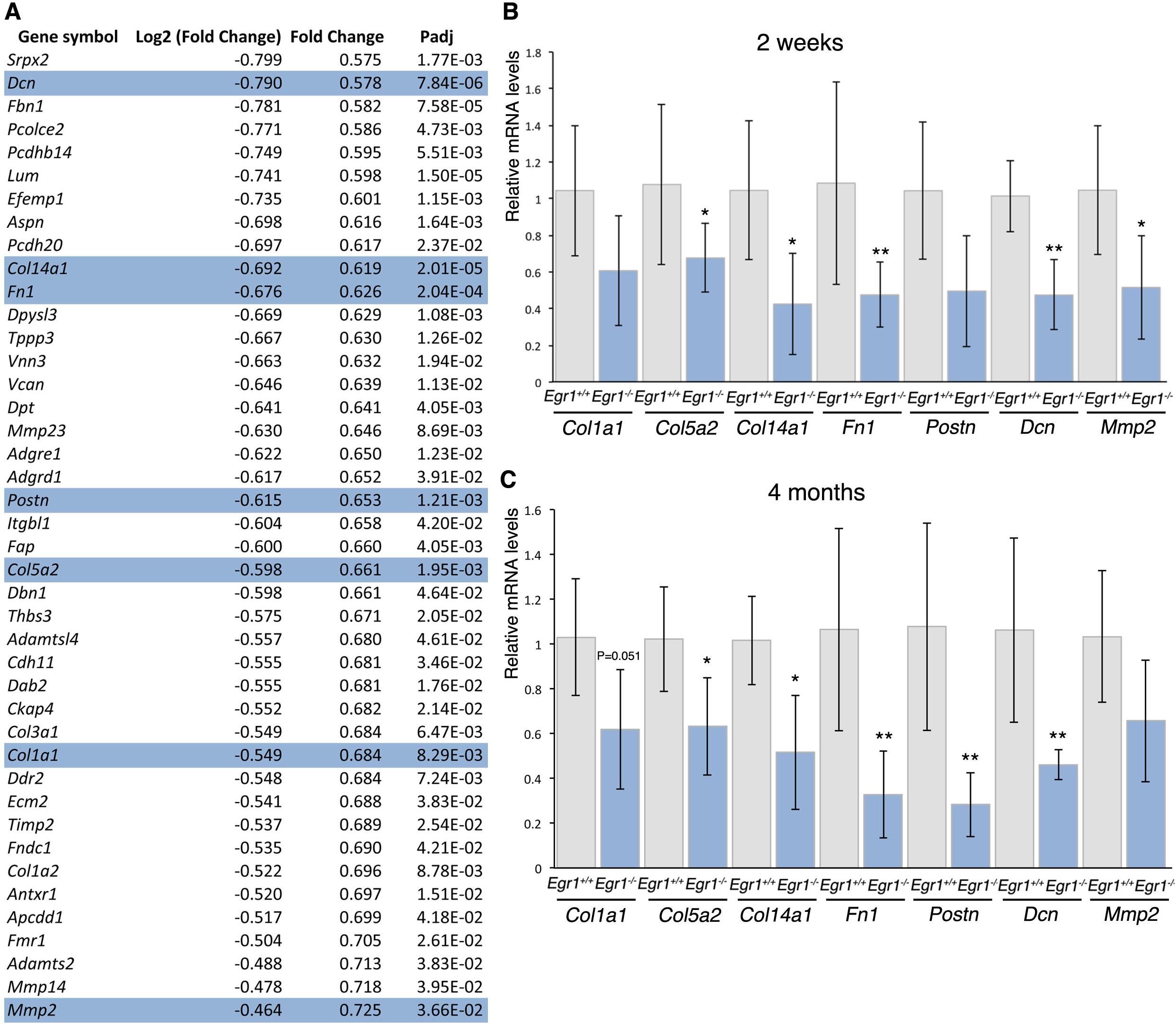
Transcriptomic analysis of the subcutaneous inguinal adipose tissue of postnatal *Egr1*^*-/-*^versus *Egr1*^*+/+*^mice reveals downregulation of extracellular matrix genes. (**A**) List of downregulated extracellular matrix genes in the inguinal subcutaneous adipose tissue of 2-week-old *Egr1*^*-/-*^mice versus wild-type mice. (**B,C**) RT-qPCR analysis of gene expression levels for extracellular matrix genes, *Col1a1, Col5a2, Col14a1, Fn1, Postn, Dcn* and *Mmp2,* in SC-WAT of 2-week-old (**B**) and 4-month-old (**C**) *Egr1*^*+/+*^and *Egr1*^*-/-*^mice. For each gene, the mRNA levels of control (*Egr1*^*+/+*^) SC-WAT were normalized to 1. Graphs show means ± standard deviations of 7 samples from 2-week-old *Egr1*^*+/+*^mice, 5 samples from 2-week-old *Egr1*^*-/-*^mice and 5 samples from 4-month-old wild type and mutant mice. The p-values were obtained using the Mann-Withney test. Asterisks indicate the p-values * p<0.05, ** p<0.01.

The concomitant upregulation of beige differentiation genes and downregulation of ECM genes is a signature of WAT browning downstream of *Egr1* deletion without any external stimulation.

### Forced *Egr1* expression in mouse mesenchymal stem cells reduces beige marker expression and promotes extracellular matrix gene expression

The spontaneous WAT browning in *Egr1*^*-/-*^mice and the direct transcriptional regulation of *Ucp1* gene by EGR1 in SC-WAT suggested that EGR1 repressed beige adipocyte differentiation. EGR1 gain-of-function experiments were performed in mouse mesenchymal stem cells, C3H10T1/2 cells, cultured under beige adipocyte differentiation conditions. Consistent with the increase in the number of adipocytes in SC-WAT of *Egr1*^*-/-*^(Figure 1), we observed a decreased number of C3H10T1/2-Egr1 cells compared to C3H10T1/2 cells after 8 days of culture in the beige differentiation medium (Figure 4A,B). Under beige stimulation, C3H10T1/2 cells acquired a beige phenotype, visualized by the appearance of numerous small lipid droplets and UCP1 expression within their cytoplasm (Figure 4A). In contrast, C3H10T1/2-Egr1 cells did not express UCP1 under beige stimulation, showing that EGR1 repressed the expression of the key thermogenic beige marker (Figure 4A). Consistent with the absence of UCP1 protein (Figure 4A), *Ucp1* mRNA levels were never increased in the presence of EGR1 (Figure 4C). This fits with EGR1 recruitment to *Ucp1* promoter in SC-WAT (Figure 2D). However, small lipid droplets were still observed in C3H10T1/2-Egr1 cells, indicating that EGR1 repressed part of the beige phenotype through the repression of UCP1, but did not fully abolish the formation of lipid droplets (Figure 4A). The expression of *Cebpb* and *Ppara* genes was significantly reduced in C3H10T1/2-Egr1 cells compared to control cells as that of *Cidea*, *Ogdh, Pank1*, *Sucla2* and *Plin5* genes (Figure 4C, Figure 4-figure supplement 1E). This showed that beige differentiation and the heat-producing ability of C3H10T1/2 cells were impaired upon EGR1 overexpression. EGR1 overexpression also blocked white adipocyte differentiation in C3H10T1/2 cells (Figure 4-figure supplement 1A-D), as previously observed (Guerquin et al., 2013). The inhibition of both beige and white differentiation programs by EGR1 is to be related with the direct (*Cebpb)* and indirect transcriptional regulation of generic adipogenesis genes by EGR1 (Figure 2).

**Figure 4.**
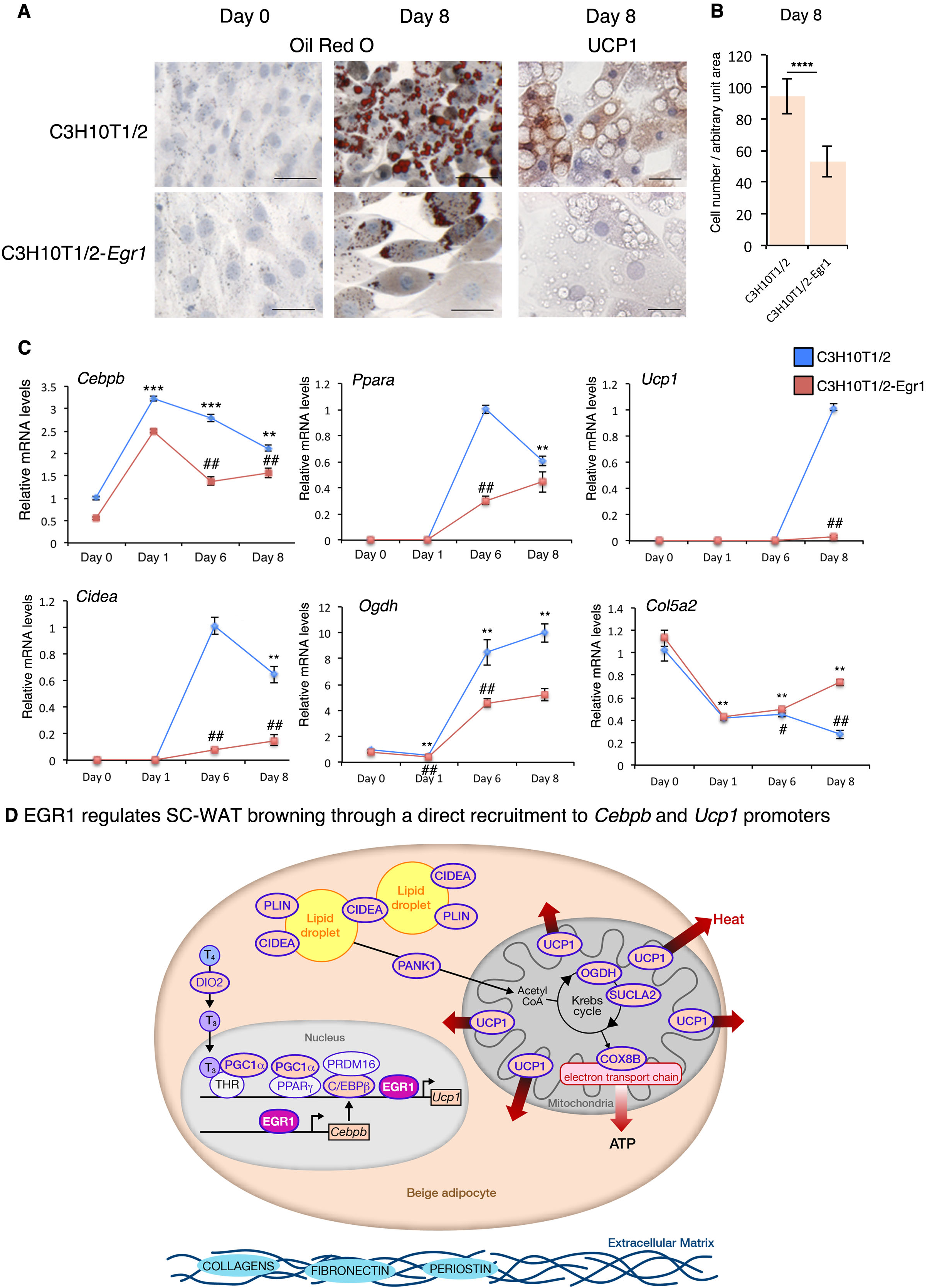
*Egr1* gain-of-function decreases beige adipocyte differentiation in mouse mesenchymal stem cells. (**A**) C3H10T1/2 and C3H10T1/2-*Egr1* cells subjected to beige adipocyte differentiation for 8 days were then stained with Oil Red O and Hematoxilin/Eosin at Day 0 (confluence) and Day 8, or immuno-stained with UCP1 antibody and counterstained with Hematoxilin/Eosin at Day 8. Scale bars: Oil red O staining 50μm, UCP1 immunostaining 25μm. (**B**) C3H10T1/2 and C3H10T1/2-Egr1 cell density after 8 days in beige differentiation medium. Graphs show means ± standard deviations of cell number from 10 pictures in each condition. The p-values were obtained using the Mann-Whitney test. Asterisks indicate the p-value **** P<0.0001. (**C**) RT-qPCR analysis of the expression levels for the adipocyte transcriptional activator *Ppara*, the beige markers, *Ucp1*, *Cidea, Plin5, Ogdh,* and the extracellular component *Col5a2* in C3H10T1/2 and C3H10T1/2-*Egr1* cells subjected to beige adipocyte differentiation. For each gene, the mRNA levels of the control C3H10T1/2 cells at Day 0 or from the first day of detection were normalised to 1. *Ogdh* and *Col5a2* expression was detected from Day 0, *Ppara*, *Cidea* and *Plin5* expression was detected from Day 6, *Ucp1* expression was detected at day 8. The graphs show the relative levels of mRNAs in C3H10T1/2 and C3H10T1/2-*Egr1* cells at different time points (Day 0, Day 1, Day 6, and Day 8) of beige adipocyte differentiation compared to C3H10T1/2 cells at Day 0 or to C3H10T1/2 cells from the first day of gene detection. For each time point, graphs show means ± standard deviations of 6 samples. The p values were calculated using the Mann-Withney test. Asterisks indicate the p-values of gene expression levels in C3H10T1/2-*Egr1* cells or C3H10T1/2 cells compared to Day 0 (*Ogdh* and *Col5a2*) or from the first day of gene detection (*Ppara*, *Cidea, Ucp1* and *Plin5)*, **P<0.01. # indicate the p-values of gene expression levels in C3H10T1/2-*Egr1* versus C3H10T1/2 cells, for each time point, ## P<0.01; # P<0.05. (**D**) Schematic cellular location of proteins encoded by *Egr1* regulated genes in SC-WAT. *Egr1* deletion upregulates the expression of genes encoding proteins involved in the beige adipocyte differentiation network (C/EBPß, PPARγ, PGC1α), thermogenesis (UCP1, COX8B, SUCLA2, OGDH), metabolism (CIDEA, PLIN5, PANK1), or thyroid hormone metabolism (DIO2). *Egr1* is expressed in adipocytes and the encoded protein is recruited to the *Cebpb* and *Ucp1* promoters. This indicates that EGR1 directly represses WAT browning. *Egr1* deletion downregulates the expression of genes encoding extracellular matrix proteins such as Collagens, Fibronectin and Perisotin.

In order to assess whether EGR1 promotes the expression of ECM genes in mesenchymal stem cells in the context of adipocyte differentiation, we analysed the expression of *Col5a2*, *Fn1* and *Postn* in C3H10T1/2 and C3H10T1/2-*Egr1* cells during beige (Figure 4C, Figure 4-figure supplement 1E) and white (Figure 4-figure supplement 1D) adipocyte differentiation. The expression of *Col5a2*, *Fn1* and *Postn* genes was upregulated in *Egr1* overexpressing cells, showing that EGR1 activated the expression of ECM genes during adipocyte differentiation. The positive regulation of ECM genes by EGR1 during adipocyte differentiation was consistent with similar regulation in the context of fibrosis, atherosclerosis and tendon repair (Guerquin et al. 2013; Buechler et al. 2015). We conclude that forced EGR1 expression in mouse mesenchymal stem cells reduces beige marker expression, while promoting ECM gene expression.

In summary, the deletion of *Egr1* induces WAT browning through recruitment to the *Cebpb* and *Ucp1* promoters in mice without any cold stimulation or fasting (Figure 5). The upregulated expression profile of beige differentiation markers and downregulated profile of ECM genes in *Egr1*-deficient WAT define a molecular signature of beige adipocyte differentiation program and constitute a protective signature against white adipocyte lipid accumulation. This study identifies *Egr1* deficiency as a therapeutic approach to counteract obesity.

## Materials and Methods

### Mouse lines

The *Egr1* gene was inactivated by homologous recombination with insertion of the *LacZ* coding sequence within the *Egr1* 5’ untranslated region in addition to a frameshift mutation upstream of the DNA-binding domain of *Egr1* (Topilko et al. 1998). The line was maintained on a C57BL/6J background (Janvier, France). All animals were kept under controlled photo-period (lights on 08:00 – 20:00 hours) and a diet of commercial rodent chow and tap water *ad libitum*. All procedures using mice were conducted in accordance with the guidelines of the French National Ethic Committee for animal experimentation N°05 and are registered under the number 01789.02.

### In situ hybridization to adipose tissue sections

Inguinal subcutaneous fat pads were isolated from 1-month-old female mice, fixed in 4% paraformaldehyde overnight and processed for in situ hybridization to 6 mm wax tissue sections as previously described (Bonnin et al., 2015). The digoxigenin-labeled mouse *Egr1* probe was used as described in Topilko et al., 1998.

### RNA isolation, sequencing and transcriptomic analysis

Fresh inguinal subcutaneous fat pads were removed from 2-week-old and 4-month-old euthanized *Egr1*^*+/+*^and *Egr1*^*-/-*^female mice and homogenized using a mechanical disruption device (Lysing Matrix A, Fast Prep MP1, 4 × 30 s, 6 m.s^-1^). Total RNA was isolated using the RNeasy mini kit (Qiagen) with 15 min of DNase I (Qiagen) treatment according to the manufacturer’s protocol. Preparation of cDNA libraries and sequencing was performed at the “Ecole Normale Supérieure” Genomic Platform (Paris, France), from subcutaneous inguinal fat pads of three 2-week-old *Egr1*^*+/+*^mice and three 2-week-old *Egr1*^*-/-*^mice. Ribosomal RNA depletion was performed with the Ribo-Zero kit (Epicentre), using 500 ng of total RNA. Libraries were prepared using the strand specific RNA-Seq library preparation ScriptSeq V2 kit (Epicentre). 51-bp paired-end reads were generated using a HiSeq 1500 device (Illumina). A mean of 56.9 ± 6.3 million reads passing the Illumina quality filter were obtained for each of the 6 samples. Reads were mapped against the *mus musculus* reference genome (UCSC Dec. 2011, GRCm38/mm10) using TopHat v2.1.0 (Kim et al., 2013), Bowtie (v2.2.5) (Langmead et al., 2012), and the Release M8 (GRCm38.p4) GTF annotations as a guide. Read counts were assigned to gene features using Feature Counts v1.4.6.p5 (Liao et al., 2014) and differential expression analysis was performed with DESeq2 v1.6.3 (Love et al., 2014). Full details of the Galaxy workflow used in this study can be retrieved via the following link: https://mississippi.snv.jussieu.fr/u/emmanuellehavis/w/copy-of-grasostendon-differential-expression-2. Gene Ontology analysis on differentially expressed genes (Padj<0.05) was performed with DAVID Bioinformatic Resources 6.8 (Huang et al., 2009). Sequencing data was uploaded to the Gene Expression Omnibus (GEO) database under the accession number GSE91058.

### Chromatin Immunoprecipitation

ChIP assays were performed with previously reported protocol (Havis et al., 2006) on the inguinal subcutaneous adipose tissue isolated from sixty 2-week-old mice, homogenized using a mechanical disruption device (Lysing Matrix A, Fast Prep MP1, 3x30 sec). Eight micrograms of the rabbit polyclonal anti-Egr-1/Krox24 (C-19) antibody (Santa Cruz Biotechnology) or 8μg of the goat anti-mouse IgG2b (Southern biotechnology) were used to immunoprecipitate 30μg of sonicated chromatin. ChIP products were analyzed by quantitative PCR. 15μg of chromatin was isolated before chromatin immunoprecipitation, to be used as positive control for the PCR experiments (Input). ChIP products and Inputs were analyzed by quantitative PCR to amplify the promoter regions upstream the *Cebpb* (-660bp;-530bp)*, Ppargc1a* (-860bp;-730bp)*, Ucp1* (-170bp; + 20bp) *and Gapdh* (-2,9Kb;-2,7Kb; negative control) coding sequences. The primer list is displayed in Supplementary table 1.

### Cell cultures

Mouse mesenchymal stem cells, C3H10T1/2 (Reznikoff et al. 1973) and the stable *Egr1* overexpressing counterparts, C3H10T1/2-Egr1 (Guerquin et al. 2013) cells, were plated on 6-well plates at a density of 33,000 cells/well and grown in Dulbecco’s Modified Eagle’s Medium (DMEM, Invitrogen) supplemented with 10% foetal bovine serum (FBS, Sigma), 1% penicillin-streptomycin (Sigma), 1% Glutamin (Sigma), 800 µg/ml G418 Geneticin (Sigma) and incubated at 37 °C in humidified atmosphere with 5% CO2.

Confluent cells were cultured in beige differentiation induction medium for 2 days and in beige maturation medium for 6 days according to published protocols (Lone et al., 2015). Day 0 corresponds to the addition of beige differentiation induction medium on confluent cells. Beige differentiation induction medium includes DMEM, 10% FBS, 1% penicillin-streptomycin, 10 *μ*g/mL Insulin (Sigma), 0.25 *μ*M Dexamethasone (Sigma), 0.5 mM 3-Isobutyl-1-methylxanthine (IBMX, Sigma), 50 nM 3.3’,5-Triiodo-L-thyronine sodium salt (T3, Sigma), 20 *μ*M Curcumin (Sigma). The beige maturation medium comprises DMEM, 10% FBS, 1% penicillin-streptomycin, 10 *μ*g/mL Insulin (Sigma), 50 nM 3,3’,5-Triiodo-L-thyronine sodium salt (T_3_, Sigma), 20 *μ*M Curcumin (Sigma), 1 *μ*M Rosiglitazone (Sigma). The maturation medium was changed every 2 days. Cells subjected to beige adipocyte differentiation medium were fixed for histological analysis or lysed for gene expression analysis at Day 0, Day 1, Day 6 and Day 8.

Confluent cells were cultured in white differentiation induction medium for 2 days and in white maturation medium for 8 days. Day 0 corresponds to the addition of white differentiation medium. White differentiation induction medium includes DMEM, 10% FBS, 1% penicillin-streptomycin, 10 *μ*g/mL Insulin (Sigma), 0.25 *μ*M Dexamethasone (Sigma), 0.5 mM 3-Isobutyl-1-methylxanthine (IBMX, Sigma), 30 nM 3.3’,5-Triiodo-L-thyronine sodium salt (T_3_, Sigma). The white maturation medium comprises DMEM, 10% FBS, 1% penicillin-streptomycin and 10 *μ*g/mL Insulin (Sigma). The maturation medium was changed every 2 days. Cells subjected to white adipocyte differentiation medium were stopped at Day 0, Day 1, Day 4 and Day 10 for histological and gene expression analysis.

Cell number measurements were performed using the free software Image J (Rasband, W.S., Image J, U. S. National Institutes of Health, Bethesda, Maryland, USA, http://imagej.nih.gov/ij/, 1997-2012).

### Oil Red O staining

C3H10T1/2 and C3H10T1/2-Egr1 cells were cultured in beige or white adipocyte differentiation medium for 8 and 10 days, respectively. Cells were fixed with 4% Paraformaldehyde (Sigma) for 15 min and washed twice with excess distilled H_2_O (Millipore). 60% Isopropanol was added for 5 min and replaced with an Oil Red O (Sigma) staining mixture, consisting of Oil Red O solution (0.5% Oil Red O dye in Isopropanol) and water in a 6:4 ratio, for 15 min. Cells were rinsed three times in distilled H_2_O, followed by a standard Hematoxylin & Eosin staining protocol.

### Immunohistochemistry

Fresh inguinal subcutaneous fat pads were removed from 1 and 4 month-old euthanized *Egr1*^*+/+*^and *Egr1*^*-/-*^ female mice, fixed in 4% formaldehyde overnight at 4°C and processed for immunohistochemistry on 10 μm wax tissue sections, as previously described (Wang et al., 2010). After wax removal, heat-induced epitope retrieval was performed by incubating sections 5 min at 95°C in Glycine-HCl buffer (0.05M Glycine, pH3.5). UCP1 protein was detected using rabbit polyclonal antibody (1:200, ab10983, Abcam), followed by secondary anti-rabbit HRP conjugate antibody (1:200, 170-6515, Biorad) and DiaminoBenzidine Tetra-Hydrochloride protocol (DAB) staining. Hematoxylin & Eosin (H&E) histological staining was performed using a standard protocol. Cell number measurements were performed using the free software Image J (Rasband, W.S., Image J, U. S. National Institutes of Health, Bethesda, Maryland, USA, http://imagej.nih.gov/ij/, 1997-2012).

C3H10T1/2 and C3H10T1/2-Egr1 cells were cultured in beige or white adipocyte differentiation medium for 8 and 10 days, respectively, on cover slips. Cells were fixed with 4% Paraformaldehyde (Sigma) for 15 min. UCP1 protein was detected using rabbit polyclonal antibody (1:200, ab10983, Abcam), followed by secondary anti-rabbit HRP conjugate antibody (1:200, 170-6515, Biorad) and DiaminoBenzidine Tetra-Hydrochloride protocol (DAB) staining. Hematoxylin & Eosin (H & E) histological staining was performed using a standard protocol. Cell number measurements were performed using the free software Image J (Rasband, W.S., Image J, U. S. National Institutes of Health, Bethesda, Maryland, USA, http://imagej.nih.gov/ij/, 1997-2012).

### Reverse-Transcription and quantitative real time PCR

For RT-qPCR analyses, 500 ng RNAs were Reverse Transcribed using the High Capacity Retrotranscription kit (Applied Biosystems). Quantitative PCR was performed using SYBR Green PCR Master Mix (Applied Biosystems) using primers listed in Supplementary Table 1.The relative mRNA levels were calculated using the 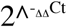 method (Livak and Schmittgen, 2001). The Cts were obtained from Ct normalized to *Rplp0*, *Rn18S* or *Actb* levels in each sample. For mRNA level analysis in SC-WAT, 5 to 7 independent RNA samples of 2-week-old and 4-month-old *Egr1*^*+/+*^and *Egr1*^*-/-*^ female mice were analysed in duplicate. For mRNA level analysis of C3H10T1/2 and C3H10T1/2-Egr1 cell cultures, 6 independent RNA samples were analysed in duplicate for each time point.

### Statistical analyses

Data was analysed using the non-parametric Mann-Withney test or ANOVA test with Graphpad Prism V6. Results are shown as means ± standard deviations. The p-values are indicated either with the value or with * or #.

## Acknowledgements

We thank Kacey Marra, Peter Rubin and Erin Kershaw from the University of Pittsburgh Medical Center, Pittsburgh, Pennsylvania, United States for comments on the manuscript and their expertise in adipose tissue biology. We thank Estelle Hirsinger from IBPS, Paris, France for comments on the manuscript. We thank Marie-Ange Bonnin from IBPS, Paris, France for technical support. We thank Sophie Lemoine and Stéphane Le Crom, from IBENS, Paris, France and Christophe Antoniewsky from ARTbio Bioinformatics Analysis Facility, Paris, France, for the bioinformatics analyses of the RNA-sequencing. We thank Sophie Gournet for illustrations.

## Author contributions

CM, acquisition, analysis and interpretation of data. KOA, contributed to unpublished essential data, analysis and interpretation of histology data, drafting the article. MO, analysis and interpretation of bioinformatics data. FC, acquisition of RNA-sequencing data. DD, conception, design, analysis and interpretation of data, drafting the article, funding. EH, conception, design, analysis and interpretation of data, drafting the article.

---

**Figure 2-figure supplement 1.**
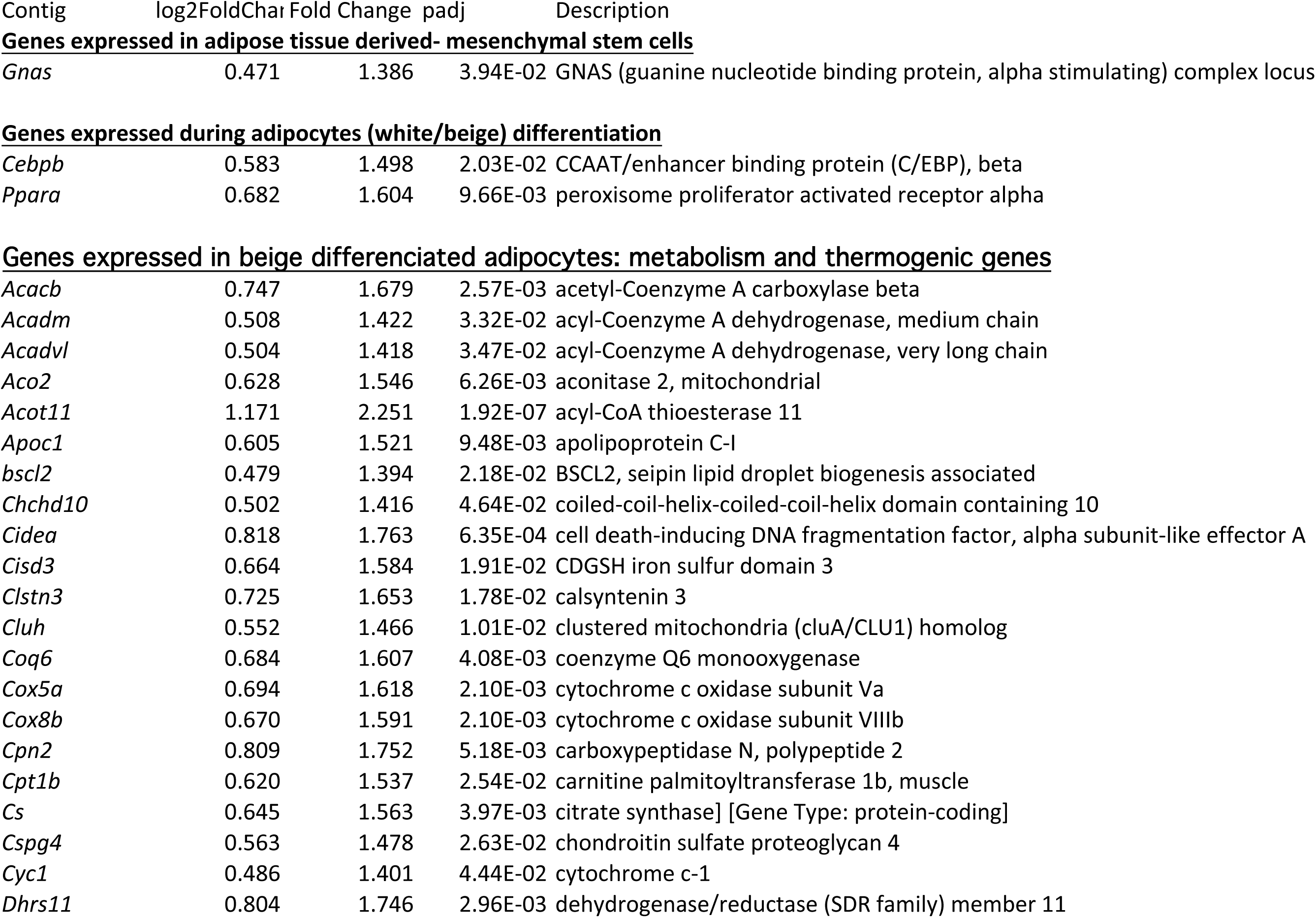
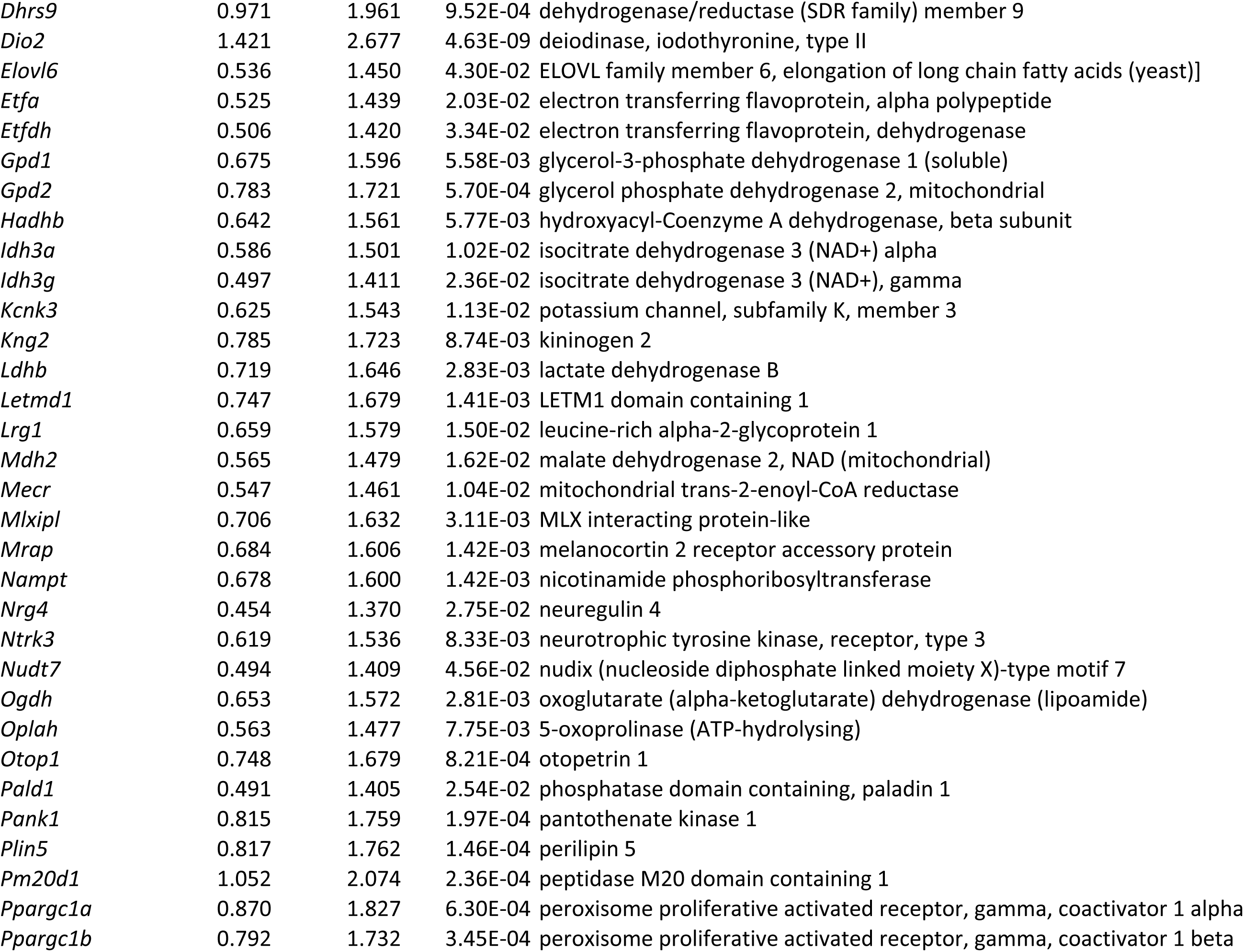
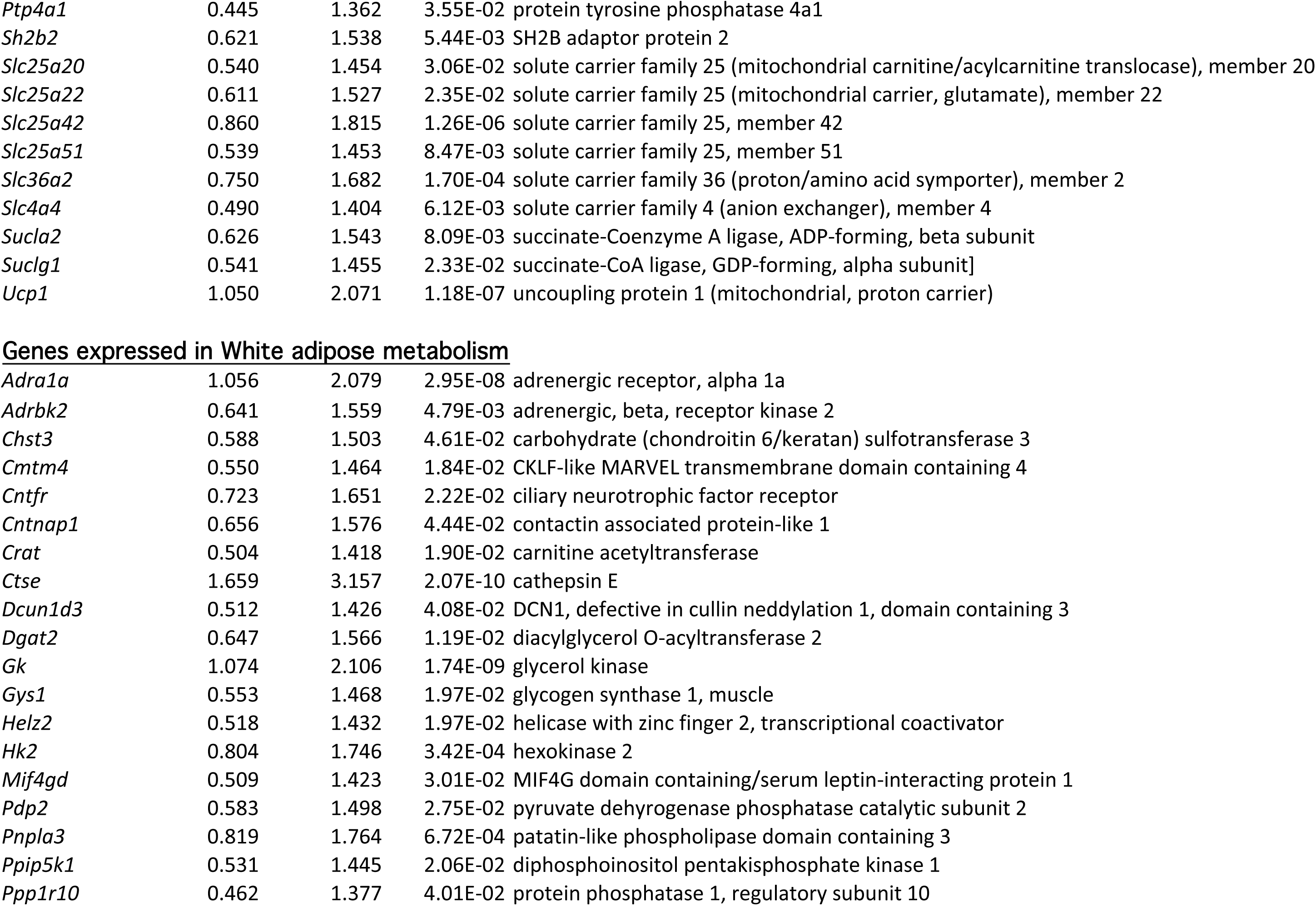
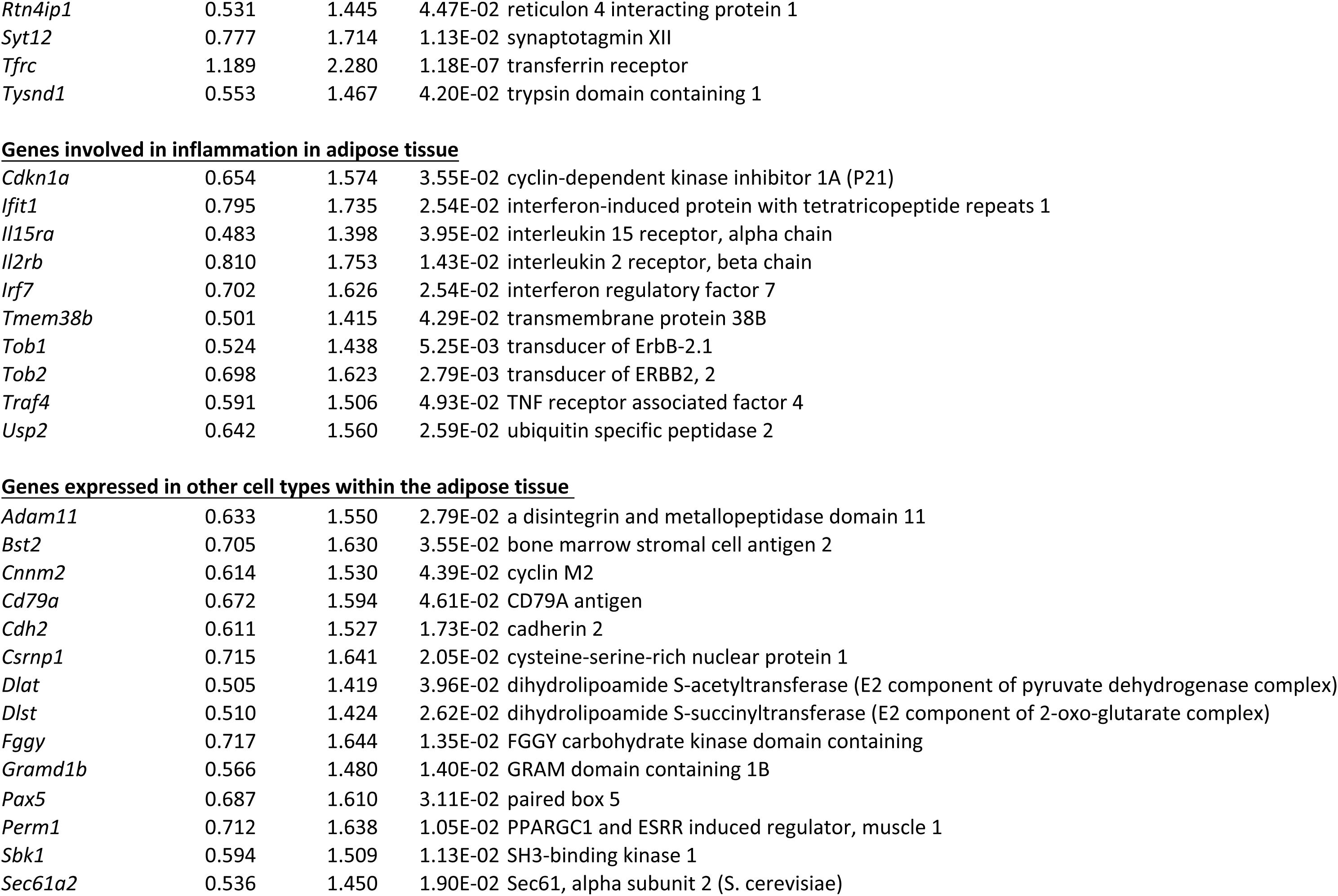
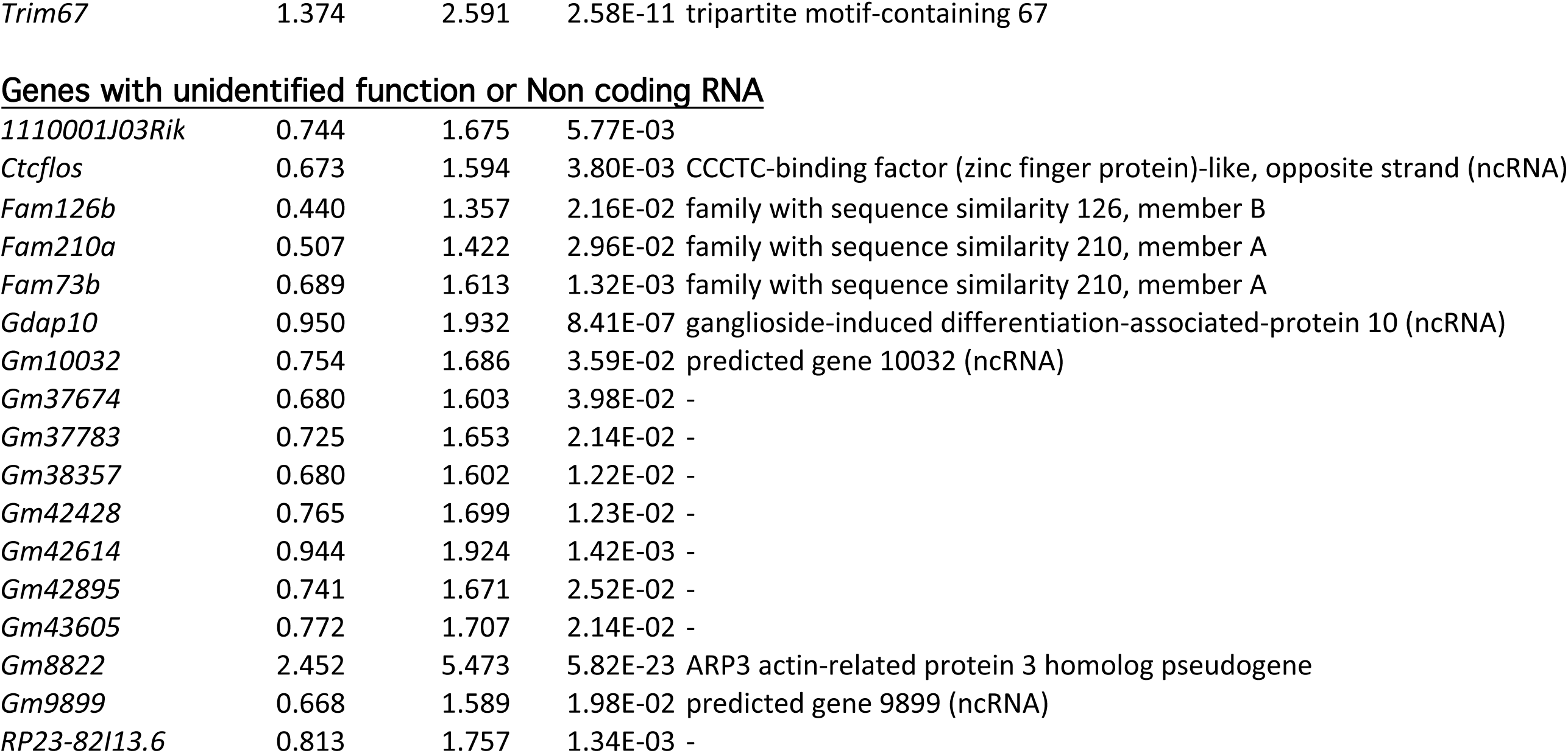
List of upregulated genes in the inguinal subcutaneous adipose tissue of 2-week-old *Egr1*^*-/-*^mice versus wild-type mice.

**Figure 2-figure supplement 2.**
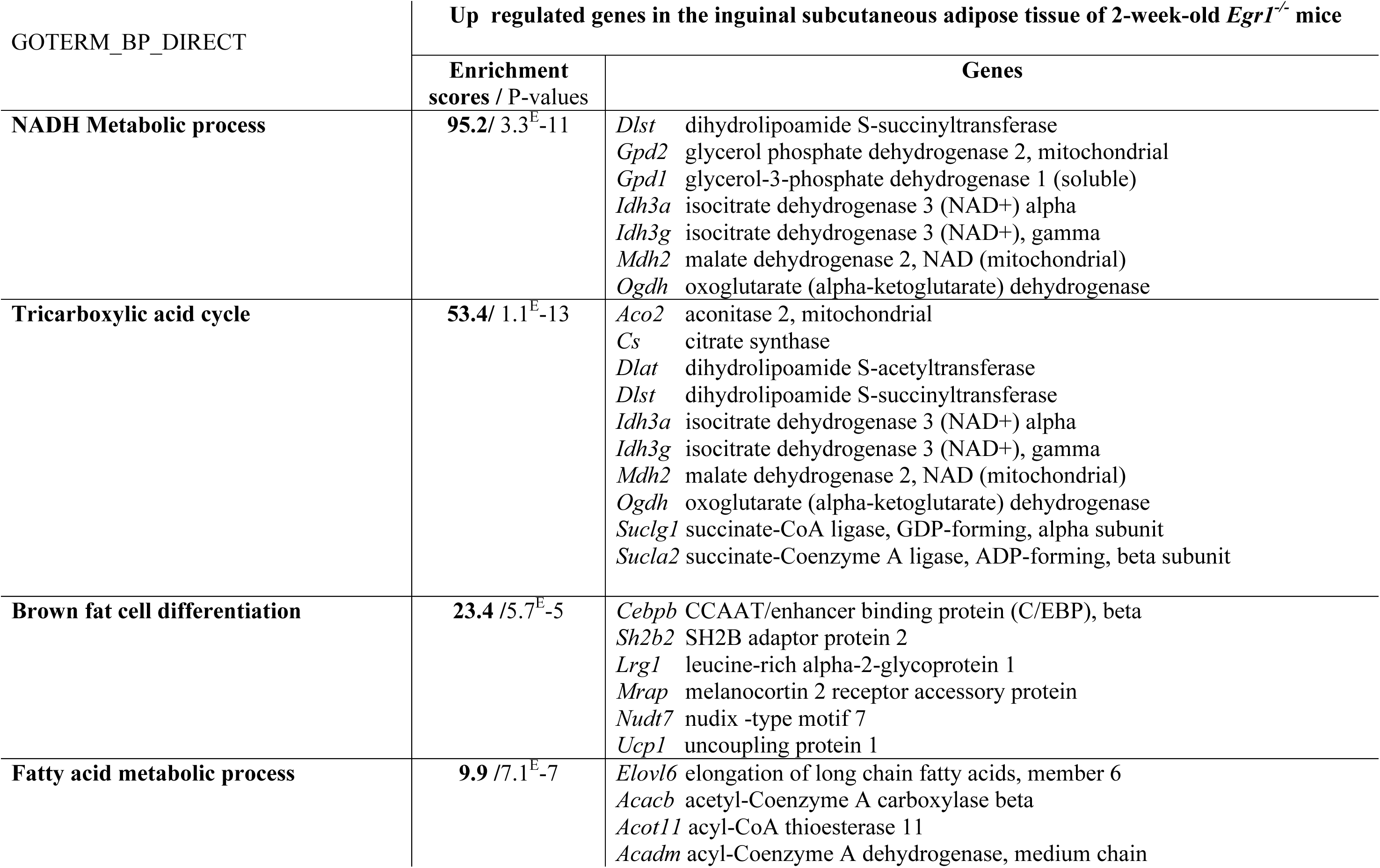
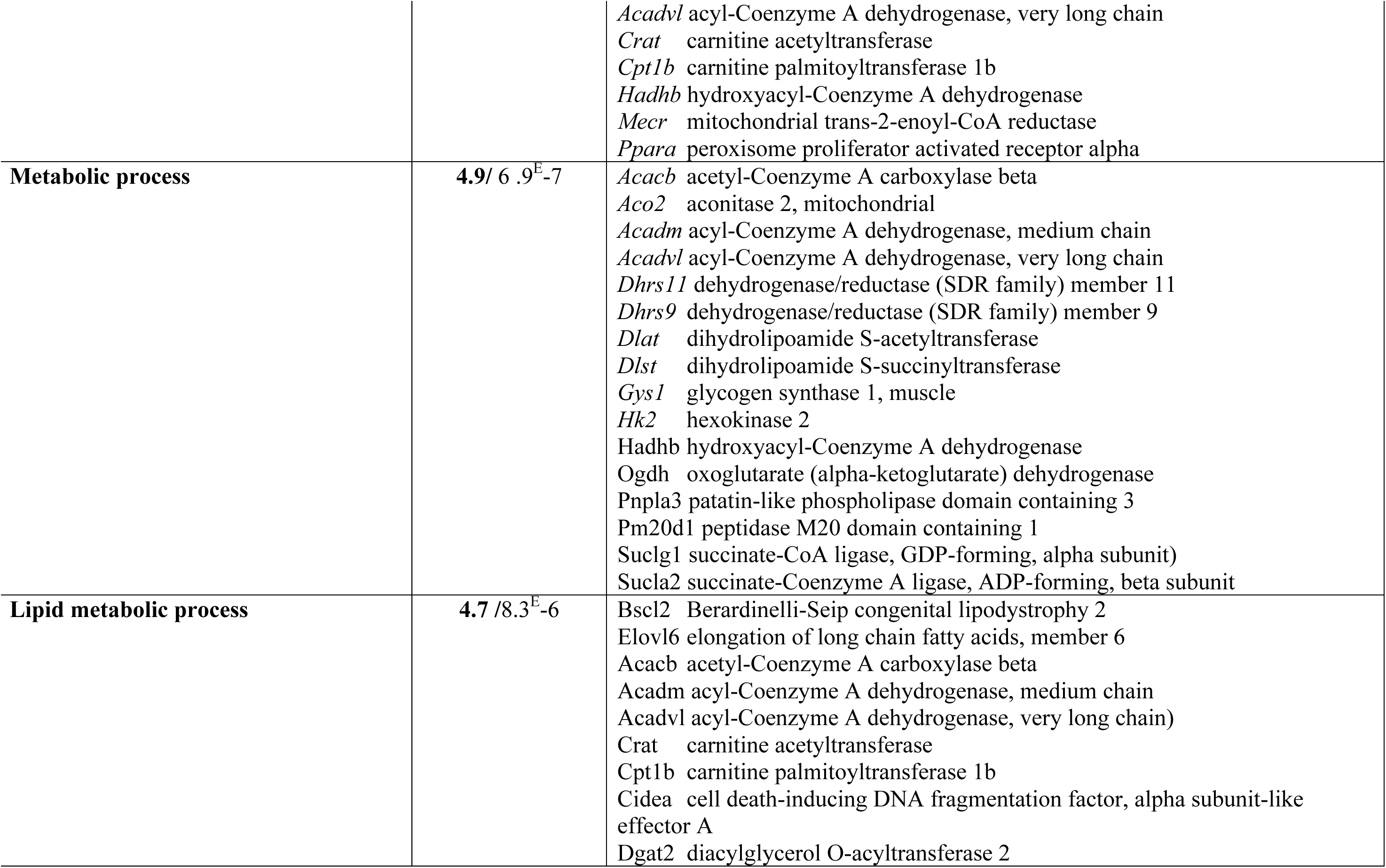
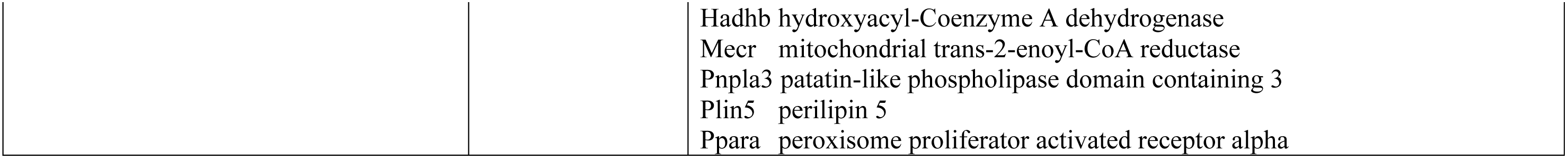
Gene Ontology analysis of the upregulated genes in the inguinal subcutaneous adipose tissue of *Egr1*^*+/+*^versus *Egr1*^*-/-*^2-week-old mice using the DAVID Bioinformatics Resources 6.8.

**Figure 3-figure supplement 1.**
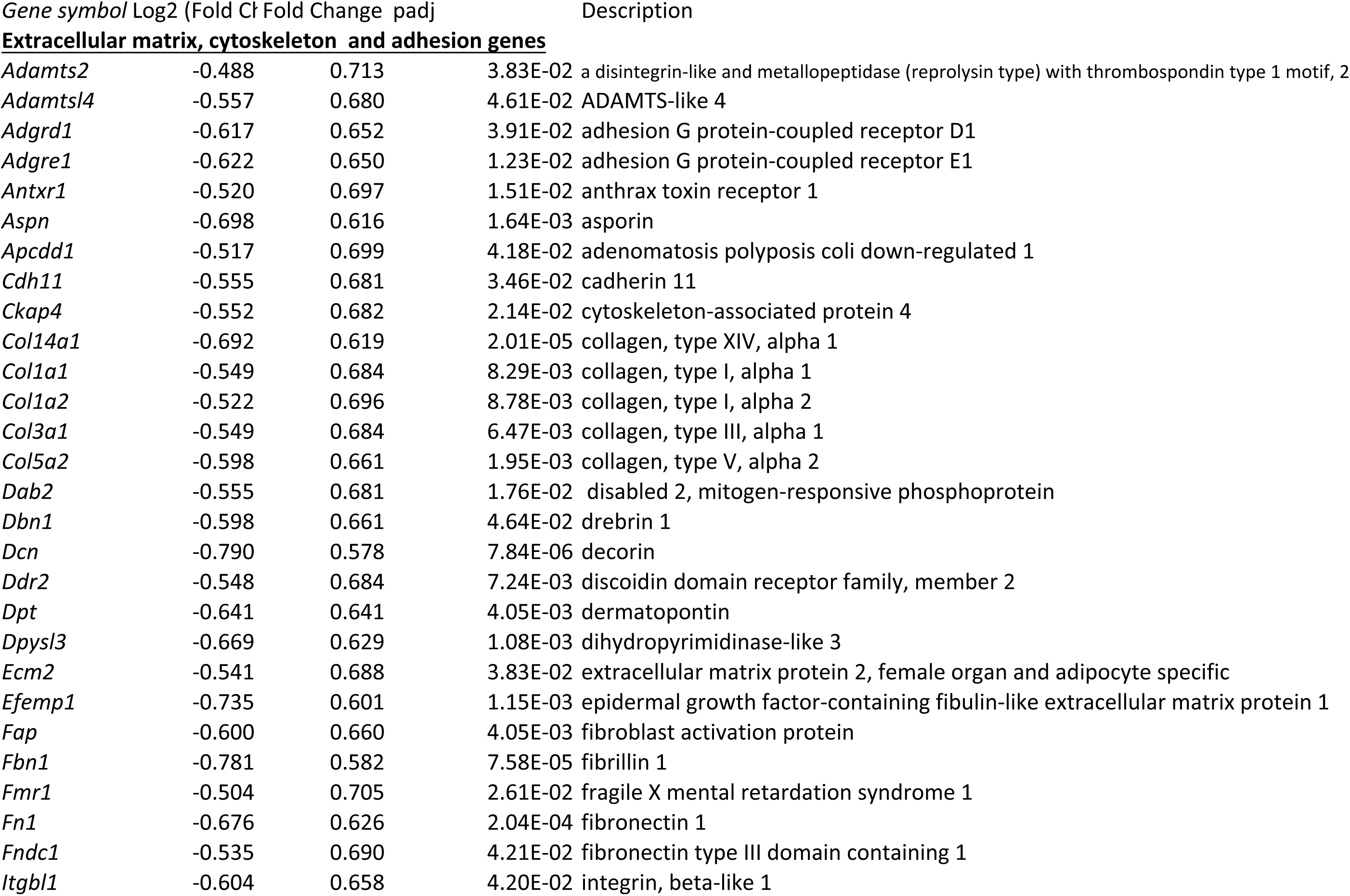
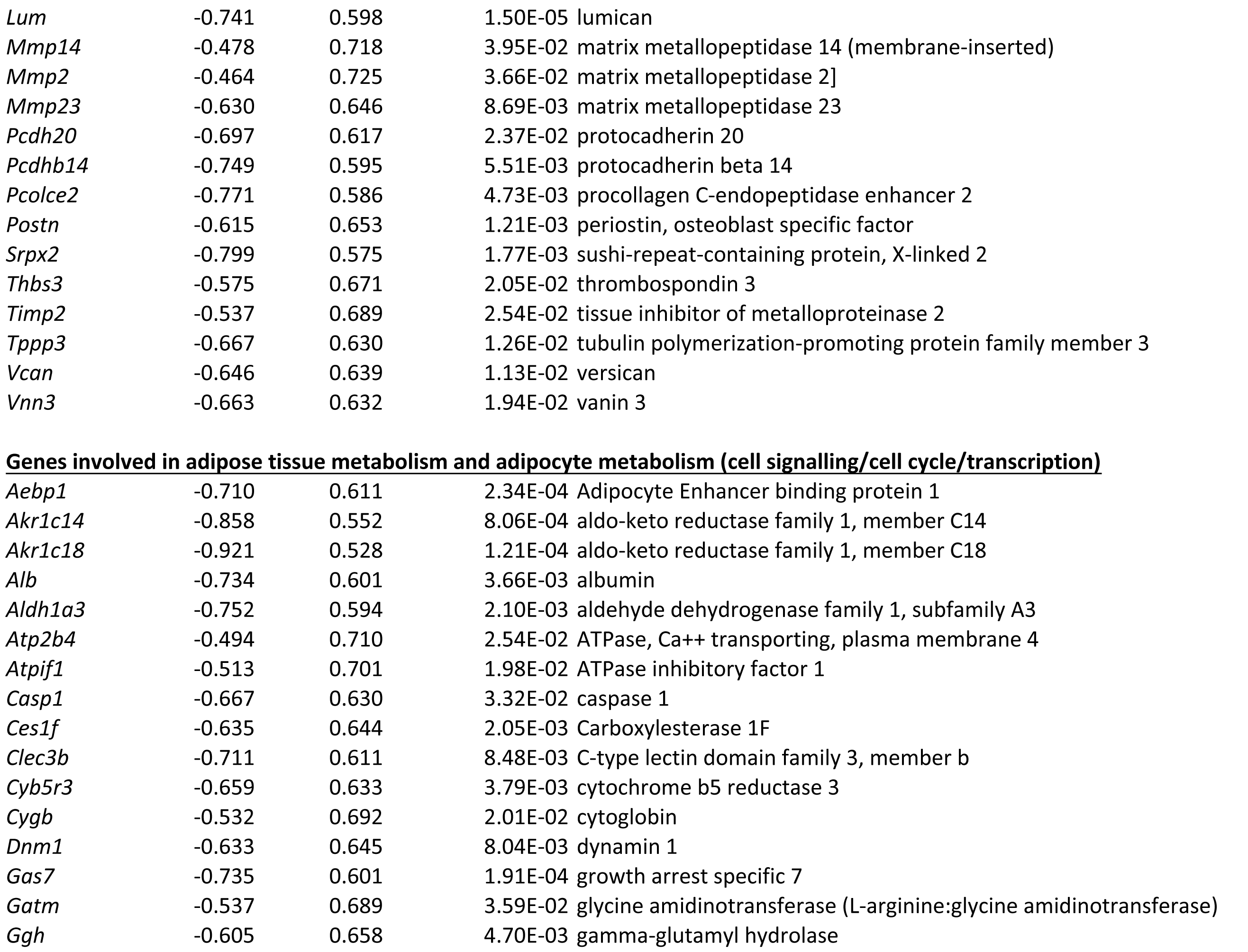
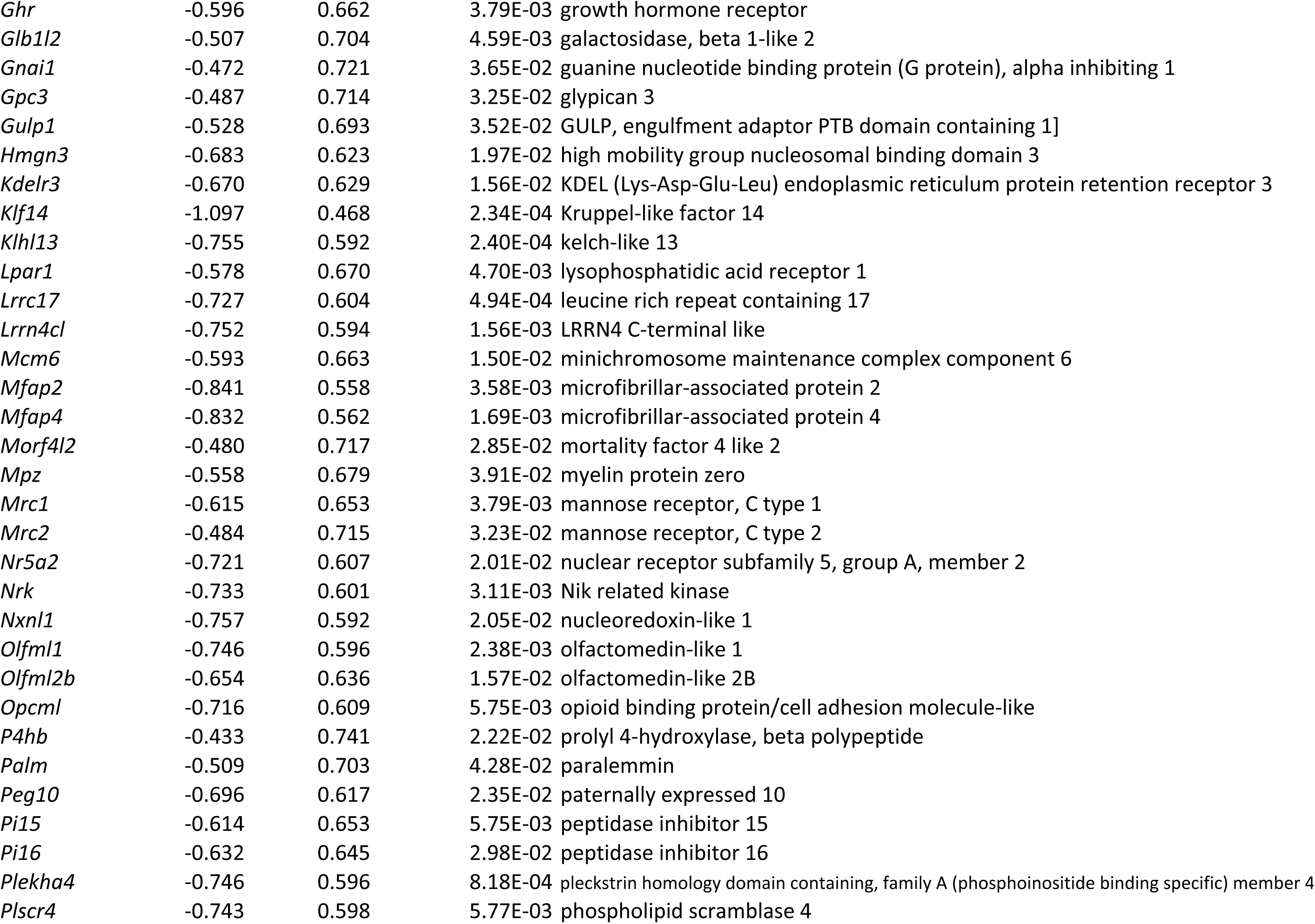
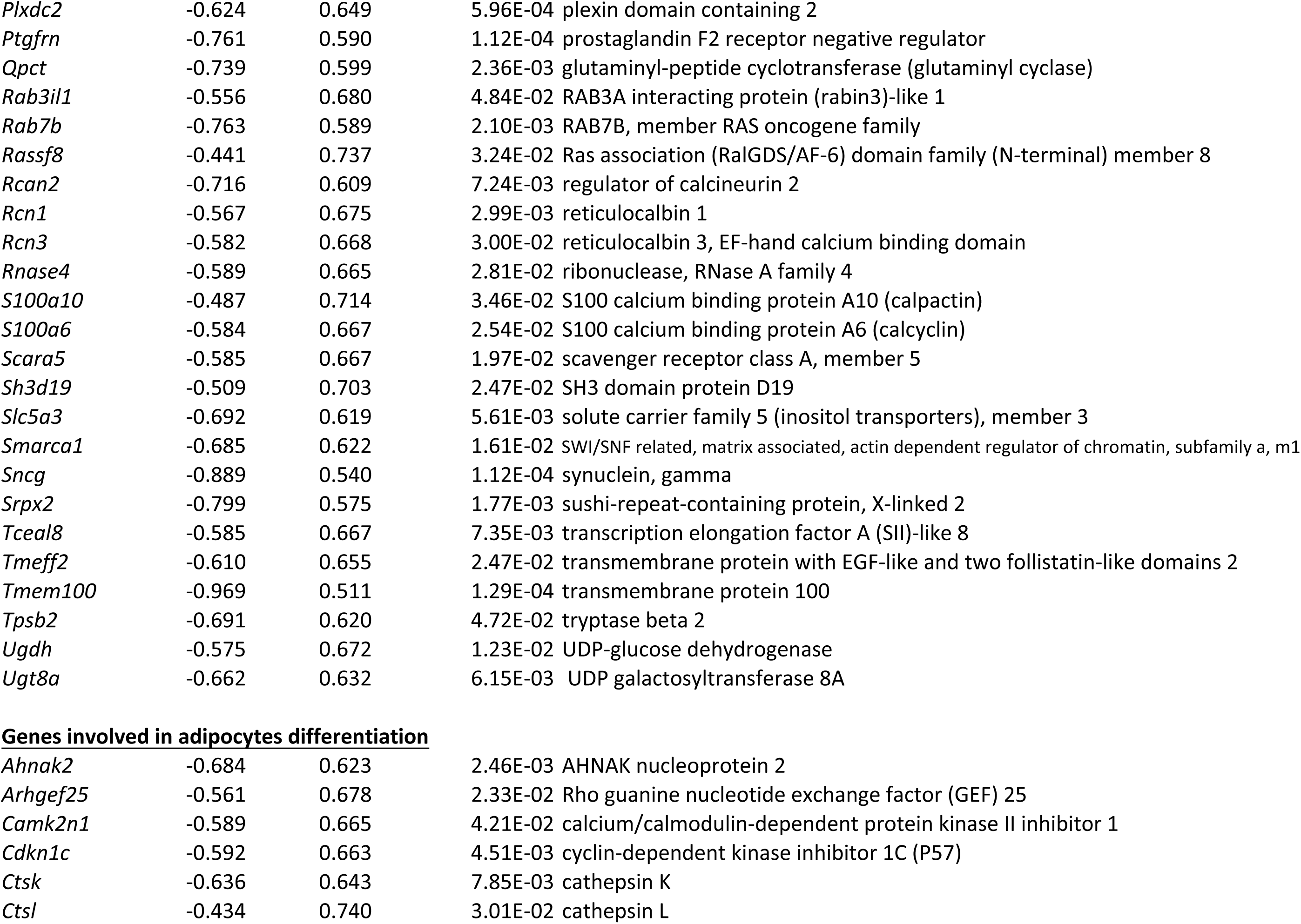
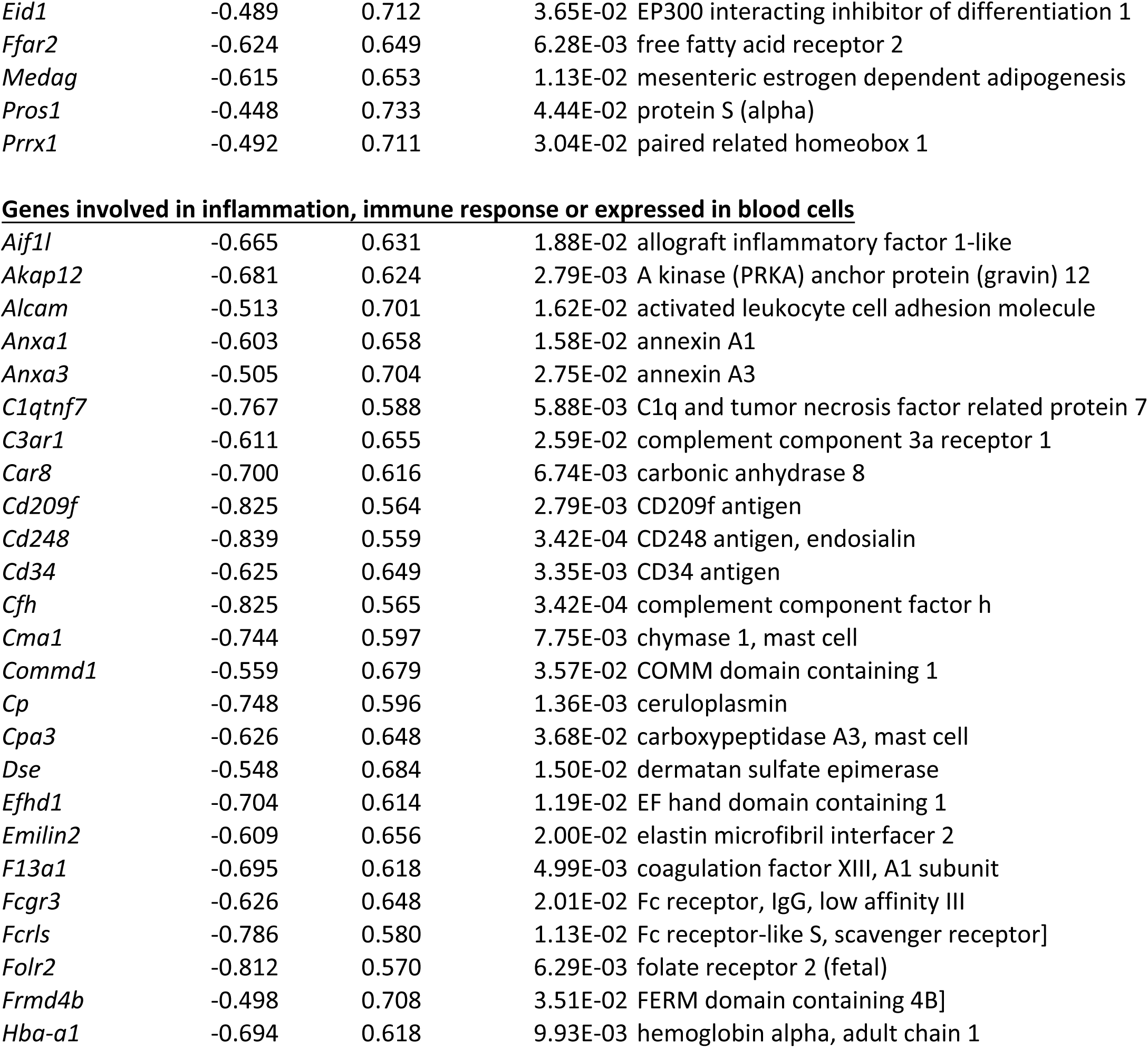
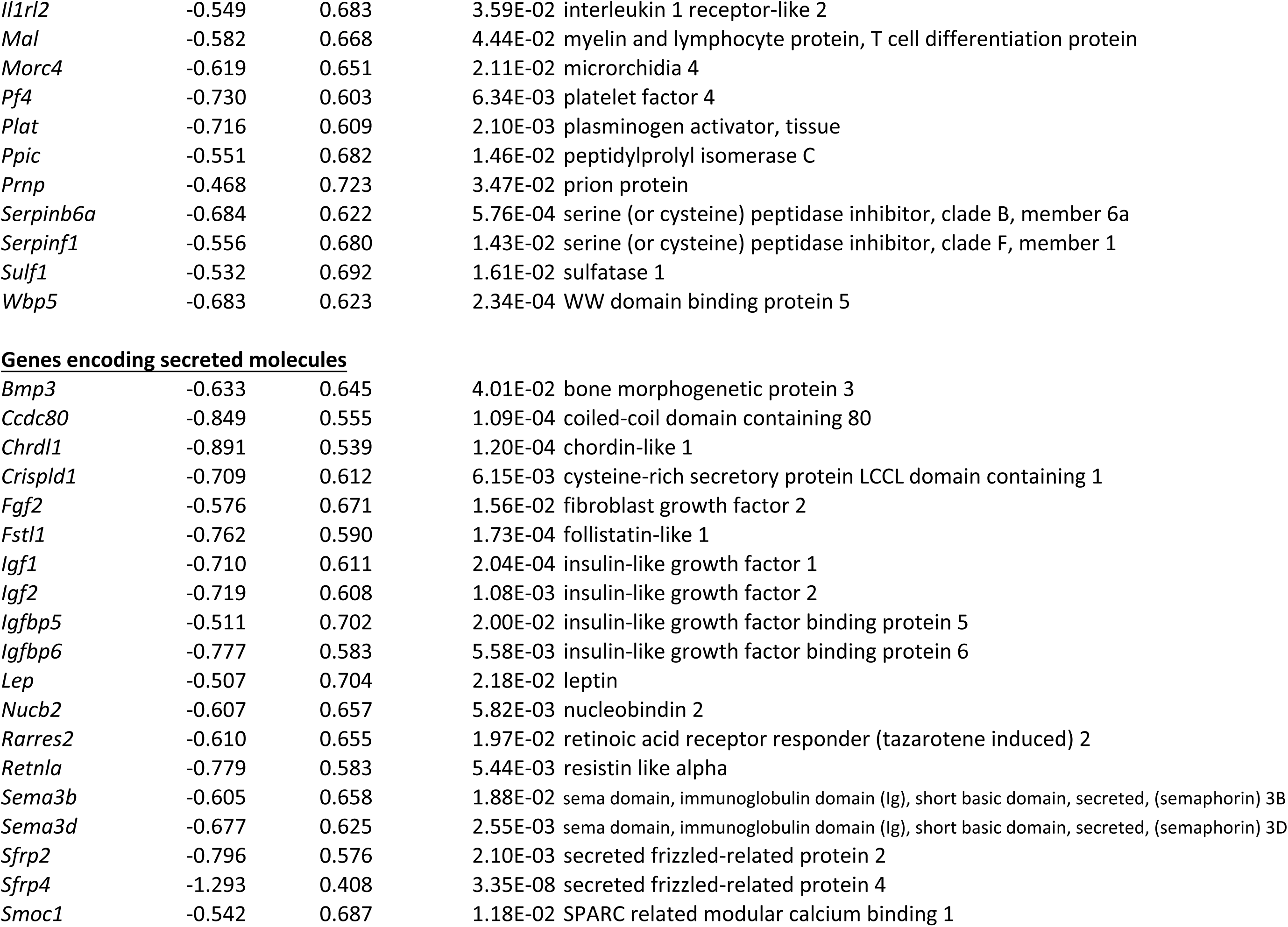
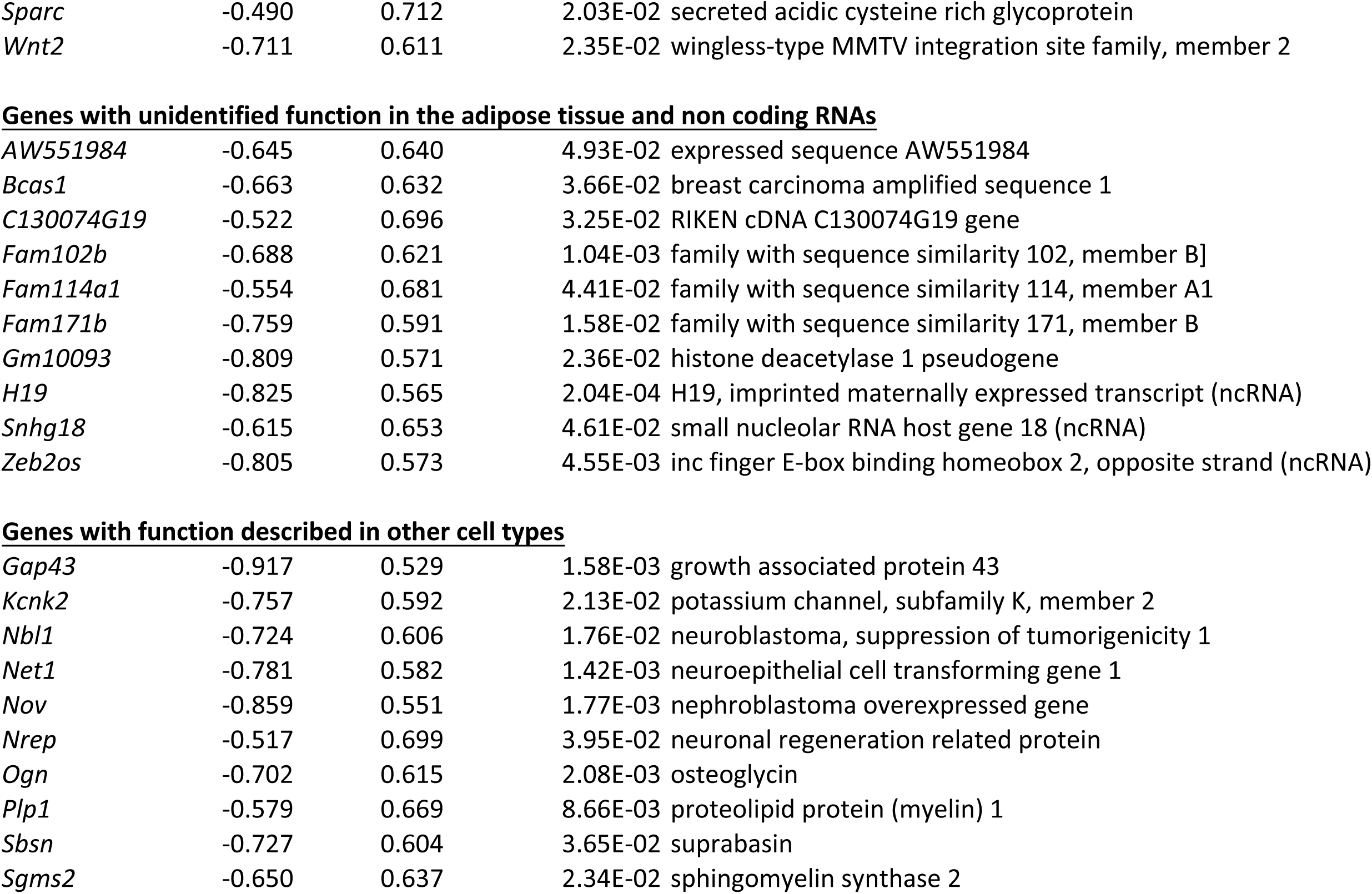
List of downregulated genes in the inguinal subcutaneous adipose tissue of 2-week – old *Egr1*^*-/-*^mice versus wild-type mice.

**Figure 3-figure supplement 2.**
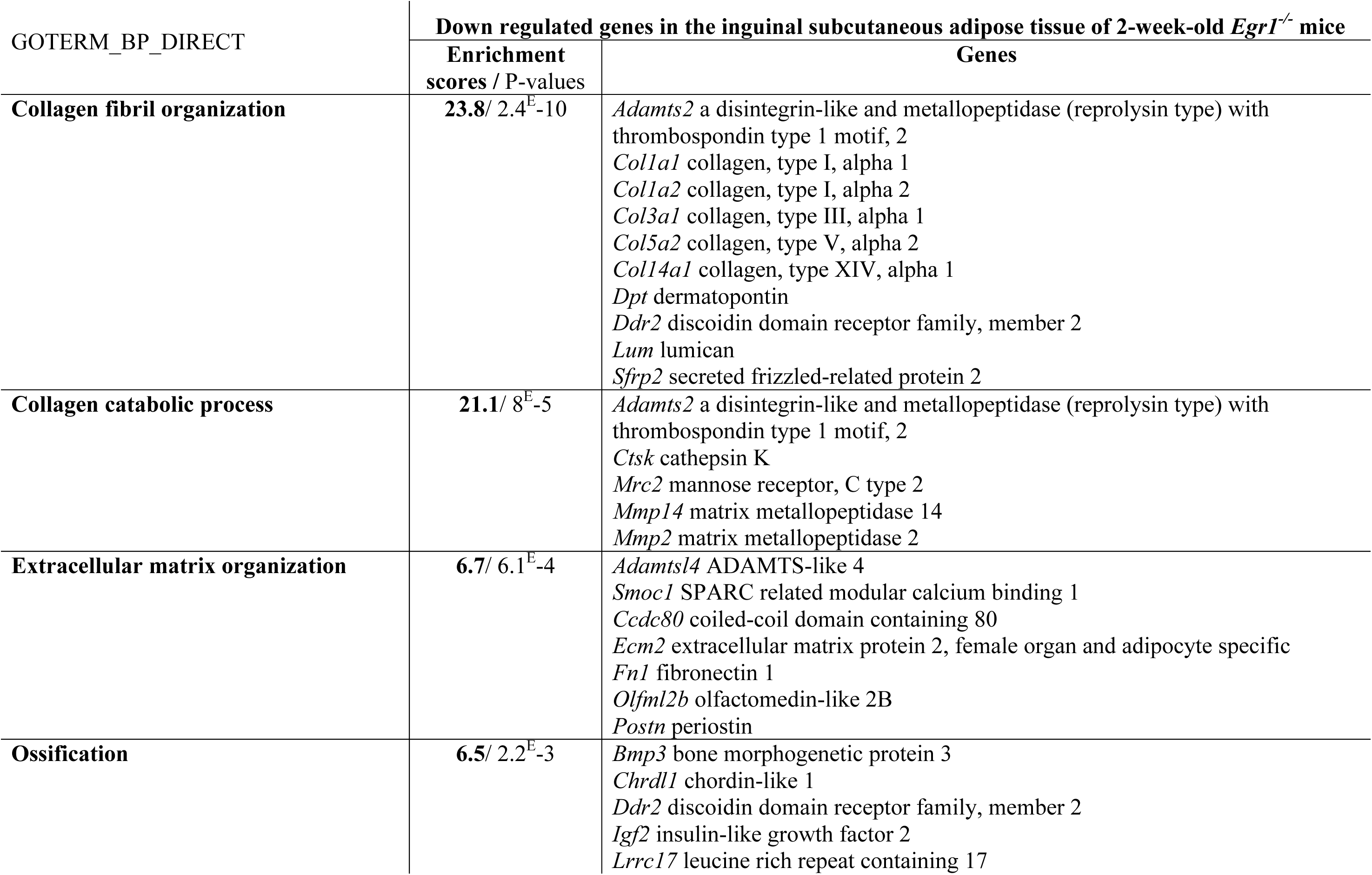
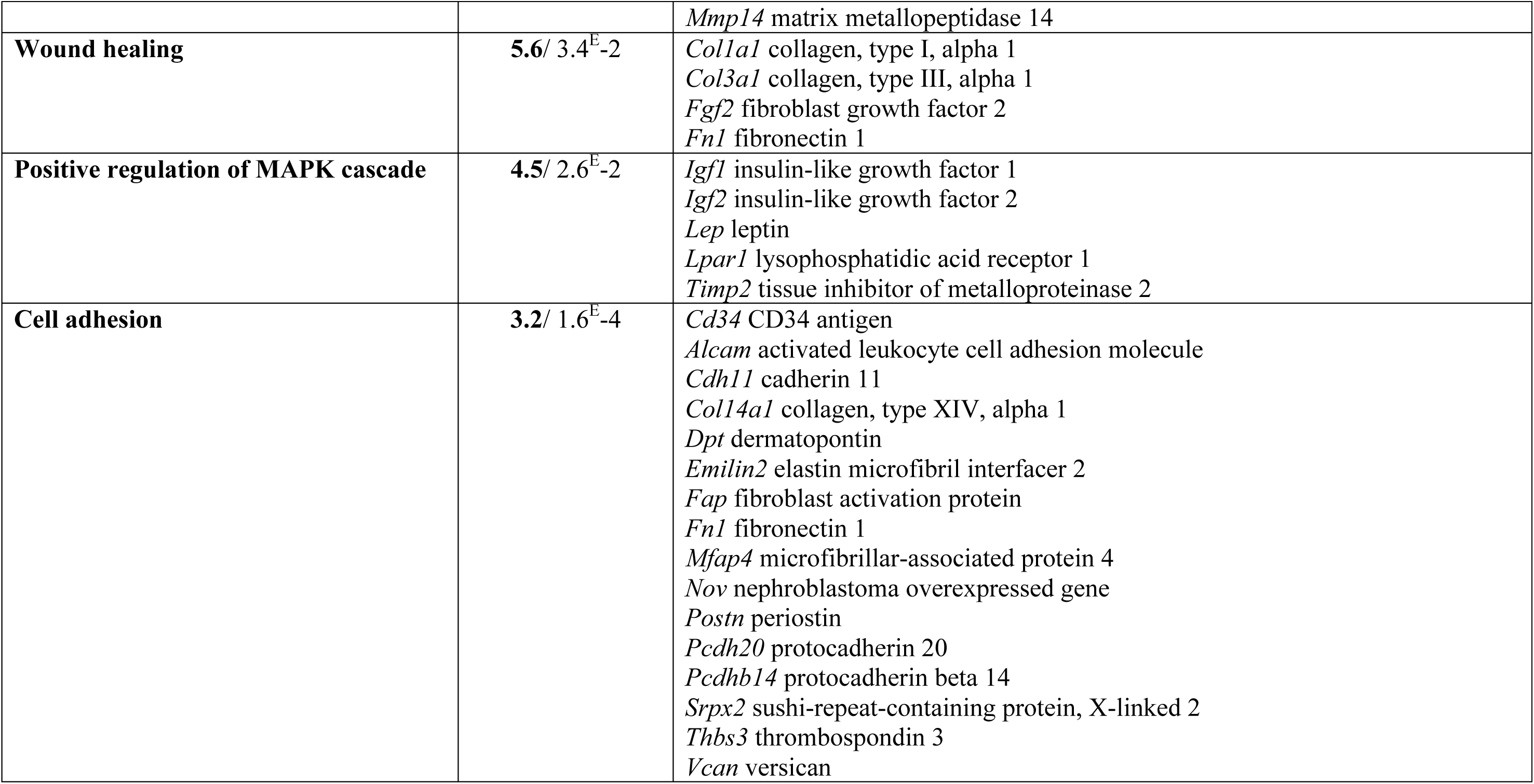
Gene Ontology analysis of downregulated genes in the inguinal subcutaneous adipose tissue of *Egr1*^*+/+*^versus *Egr1*^*-/-*^2-week-old mice using the DAVID Bioinformatics Resources 6.8.

**Figure 4-figure supplement 1.**
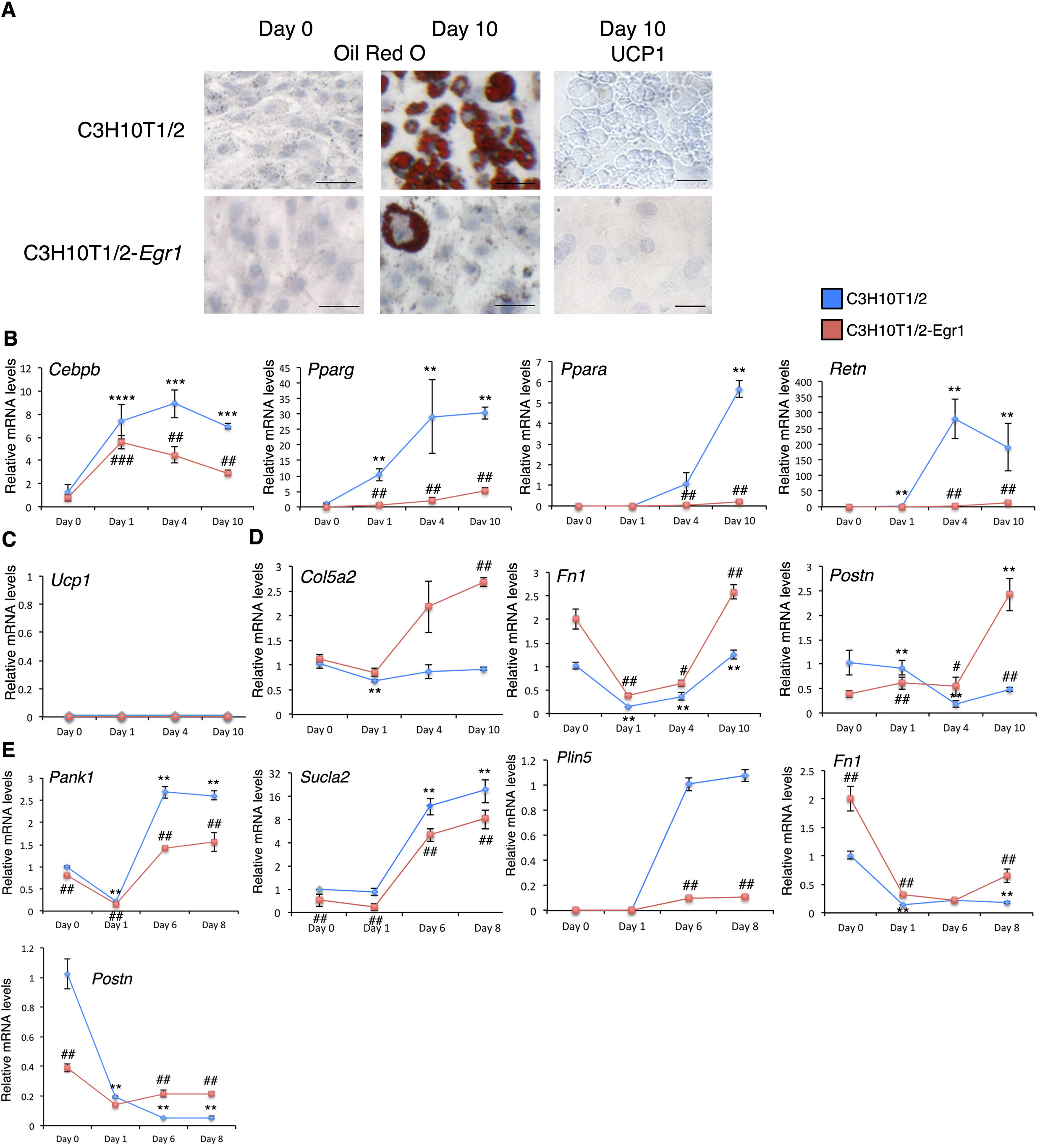
*Egr1* gain-of-function decreases white and beige adipose tissue differentiation in mouse mesenchymal stem cells. (**A**) C3H10T1/2 and C3H10T1/2-*Egr1* cells subjected to white adipocyte differentiation for 10 days were stained with Oil Red O and Hematoxilin/Eosin at Day 0 (confluence) and Day 10, or immuno-stained with UCP1 antibody and counterstained with Hematoxilin/Eosin at Day 10. UCP1 was never found to be expressed in cells cultured in white adipocyte differentiation medium. Scale bars: Oil red O staining 50μm, UCP1 immunostaining 25μm. (**B-E**) RT-qPCR analysis of the expression levels for the generic adipocyte differentiation genes *Cebpb, Pparag, Ppara*, the white differentiation marker *Retn* (**B**), the thermogenic marker *Ucp1* (**C**), the extracellular matrix genes *Col5a2*, *Fn1* and *Postn* (**D**), in C3H10T1/2 and C3H10T1/2-*Egr1* cells subjected to 10 days of white adipocyte differentiation conditions. *Egr1* repressed the expression of *Cepbb*, *Pparg Ppara* and *Retn*, involved in the white adipocyte differentiation program and activated the expression of ECM genes, *Col5a2*, *Fn1* and *Postn* during white adipocyte differentiation. *Ucp1* expression was not detected in cells cultured in white differentiation conditions. (**E**) RT-qPCR analysis of the expression levels for the generic adipocyte differentiation marker *Cebpb*, beige adipocyte markers *Pank1 and Sucla2* and ECM genes *Fn1* and *Postn* in C3H10T1/2 and C3H10T1/2-*Egr1* cells subjected to 8 days of beige adipocyte differentiation. EGR1 repressed the expression of *Cepbb* and the beige adipocyte markers *Pank1 and Sucla2.* EGR1 activated the expression of ECM genes *Col5a2*, *Fn1* and *Postn* during beige adipocyte differentiation. The mRNA levels of the C3H10T1/2 cells at day 0 or from the first day of detection were normalized to 1, so the graphs show the relative levels of mRNA in the C3H10T1/2 (n = 6) and C3H10T1/2-*Egr1* cells (n = 6) at different time points (Day 0, Day 1, Day 4, and Day 10 of white adipocyte differentiation or Day 0, Day 1, Day 6 and Day 8 of beige adipocyte differentiation) compared to C3H10T1/2 cells at day 0 or from the first day of gene detection. Error bars indicate standard deviations. The p values were calculated using the Mann-Withney test. Asterisks indicate the p-values of gene expression levels in C3H10T1/2-*Egr1* cells or C3H10T1/2 cells compared to Day 0 (*Cebpb, Pparg, Col5a2, Fn1, Postn, Pank1, Sucla2*) or from the first day of gene detection (*Ppara*, Retn), *** P<0.001, **P<0.01. # indicate the p-values of gene expression levels in C3H10T1/2-*Egr1* versus C3H10T1/2 cells, for each time point, ## P<0.01; # P<0.05.

**Supplementary Table 1.**
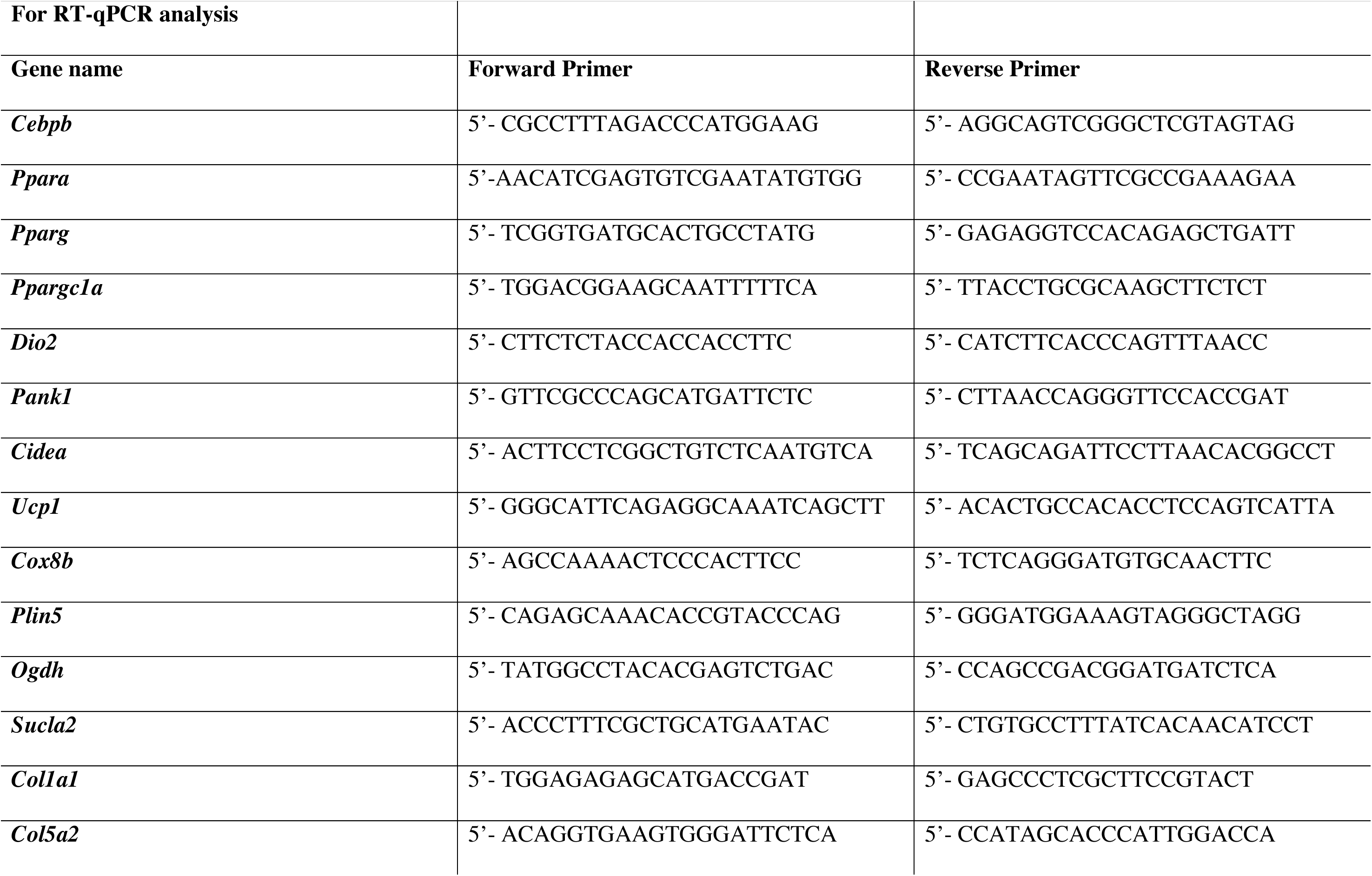
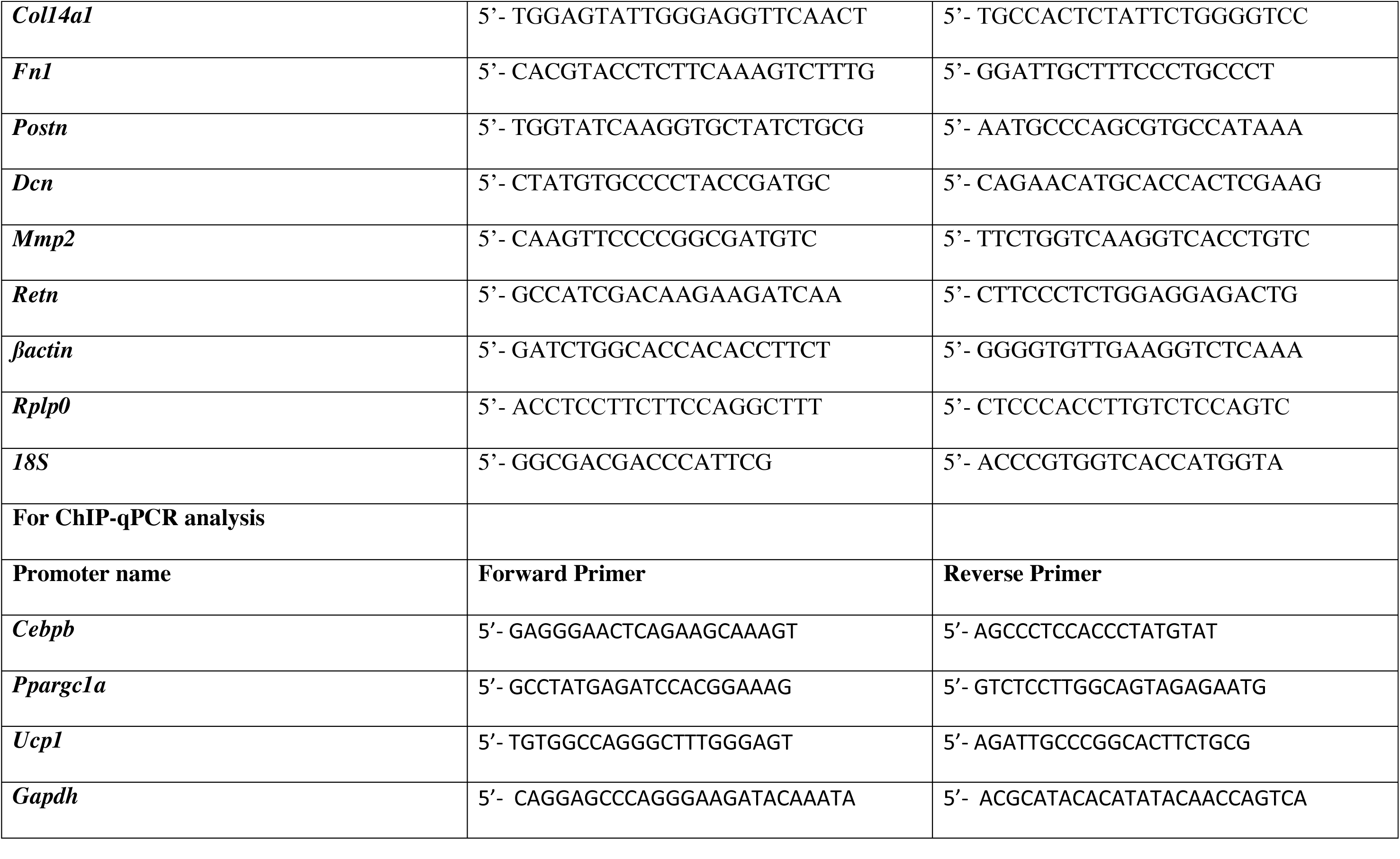
List of primers used for quantitative PCR.

